# The roles of Amh in zebrafish gonad development and sex determination

**DOI:** 10.1101/650218

**Authors:** Yi-Lin Yan, Peter Batzel, Tom Titus, Jason Sydes, Thomas Desvignes, Ruth Bremiller, Bruce Draper, John H. Postlethwait

## Abstract

Fetal mammalian testes secrete Amh (Anti-Müllerian hormone), which inhibits female reproductive tract (Müllerian duct) development. Amh also derives from mature mammalian ovarian follicles, which marks oocyte reserve and characterizes PCOS (polycystic ovarian syndrome). Zebrafish (*Danio rerio*) lacks Müllerian ducts and the Amh receptor gene *amhr2* but, curiously, retains *amh*. To discover the roles of Amh in the absence of Müllerian ducts and the ancestral receptor gene, we made *amh* null alleles in zebrafish. Results showed that normal *amh* prevents female-biased sex ratios. Adult male *amh* mutants had enormous testes, half of which contained immature oocytes, demonstrating that Amh regulates male germ cell accumulation and inhibits oocyte development or survival. Mutant males formed sperm ducts and some produced a few offspring. Young female mutants laid a few fertile eggs, so they also had functional sex ducts. Older *amh* mutants accumulated non-vitellogenic follicles in exceedingly large but sterile ovaries, showing that Amh helps control ovarian follicle maturation and proliferation. RNA-seq data partitioned 21-day post-fertilization (dpf) juveniles into two groups that each contained mutant and wild type fish. Group21-1 up-regulated ovary genes compared to Group21-2, which were likely developing as males. By 35dpf, transcriptomes distinguished males from females and, within each sex, mutants from wild types. In adult mutants, ovaries greatly under-expressed granulosa and theca genes and testes under-expressed Leydig cell genes. These results show that ancestral Amh functions included development of the gonadal soma in ovaries and testes and regulation of gamete proliferation and maturation. A major gap in our understanding is the identity of the gene encoding a zebrafish Amh receptor; we show here that the loss of *amhr2* is associated with the breakpoint of a chromosome rearrangement shared among cyprinid fishes.

**Article Summary:** Anti-Müllerian hormone (Amh) inhibits female reproductive duct development, signals oocyte reserve, and marks polycystic ovarian syndrome. Zebrafish lacks Müllerian ducts and the typical Amh receptor, questioning evolving roles of Amh. We made knockout mutations in zebrafish *amh*. Most mutants were female and the few males often had oocytes in their testes, showing that Amh promotes male development. Mutant reproductive ducts functioned, but testes were enormous and ovaries accumulated immature oocytes, showing that Amh regulates germ cell proliferation and maturation. Transcriptomics revealed that Amh controls development of steroid-producing gonad cells. Amh in zebrafish preserved ancestral roles despite losing Müllerian ducts and the Amh receptor.

## INTRODUCTION

Developing mammalian embryos form the rudiments of both male and female sex ducts, the Wolffian and Müllerian ducts, respectively. Over 70 years ago, Alfred Jost conducted remarkable experiments to learn if gonads control sex duct development (Jost 1947). He removed undifferentiated gonads from rabbit fetuses and re-implanted them into the uterus of surrogate rabbit hosts. Gonadectomized kits lost male sex ducts but retained female sex ducts. He concluded that developing testes maintain male ducts (epididymis, seminal vesicles, and vas deferens) but destroy female sex duct anlagen (fallopian tubes and uterus). In contrast, developing ovaries neither maintain male ducts nor destroy female ducts. Subsequent experiments showed that one testis-derived substance (testosterone) maintains male sex duct rudiments and another (anti-Müllerian hormone, AMH, also called Müllerian Inhibiting Substance, MIS), inhibits female reproductive duct anlagen (Elger 1966; Josso 1972).

Although AMH from testes represses female duct development, AMH from ovaries begins to appear in the third trimester of human fetal development from primary and preantral follicles (Munsterberg AND Lovell-badge 1991). Ovarian *AMH* expression peaks in juvenile women, declines with age, and disappears at menopause; thus, circulating AMH levels reflect a woman’s ovarian follicle reserve (Visser *et al*. 2006; Zec *et al*. 2011). Investigations of *Amh* mutant mice showed that chromosomal XY males that lack Amh activity develop oviducts, uterus, and vagina in addition to male reproductive ducts (Behringer *et al*. 1994). Testes in Amh-deficient XY mice attain normal size, but some show Leydig cell hyperplasia (Behringer *et al*. 1994). Chromosomally female XX *Amh* mutant juvenile mice have more preantral and small antral follicles and older mutant females have fewer primordial follicles, preantral, and small antral follicles than wild-type siblings (Behringer *et al*. 1994; Durlinger *et al*. 1999), suggesting that without AMH, primordial follicles develop more rapidly than normal, which results in larger juvenile ovaries that lose follicles prematurely. This property led to the use of circulating AMH as a marker of polycystic ovarian syndrome (PCOS), the most common problem for couples who visit fertility clinics (Pigny *et al*. 2003; Diamanti-KANDARAKIS 2008)].

We wondered about the evolution of AMH functions and their relationship to reproductive ducts. Jawless fish lack specialized gamete-transporting sex ducts; lamprey gonads release gametes directly into the body cavity where they are forced out during spawning through genital pores (Applegate 1948; Hardisty 1971). Cartilaginous fish evolved paired Müllerian ducts (or paramesonephric ducts) that condense from intermediate mesoderm parallel to Wolffian ducts (mesonephric ducts), and differentiate into the female reproductive tract, including the fallopian tubes, which collect oocytes released into the coelomic cavity (Wourms 1977). Among bony fish, tetrapods and basally diverging ray-finned fish like spotted gar (Ferrara AND Irwin 2001) maintained this ancestral state, but teleosts lost their Müllerian ducts; gonoducts in many teleosts develop from somatic cells posterior to the gonad, and gametes pass from the gonad directly into the ducts rather than into the body cavity (e.g (Suzuki and Shibata 2004; Kossack *et al*. 2019)). We therefore wondered how Amh functions evolved in a teleost given that its eponymous feature of Müllerian duct inhibition is no longer relevant in the absence of a Müllerian duct.

Despite the absence of Müllerian ducts, Amh performs a reproductive function in at least some teleosts because a Y-chromosome variant of *amh* (*amhY*) plays a role in sex determination in the Patagonian pejerrey (Hattori *et al*. 2012) and a variant Amh receptor (Amhr2) acts in sex determination in several, but not all, species of pufferfish (Kamiya *et al*. 2012; Ieda *et al*. 2018). In addition, *amhr2* mutants in medaka show excess germ cell proliferation, premature male meiosis, sex reversal in some chromosomally XY fish, and early stage follicular arrest in females (Morinaga *et al*. 2007). We lack, however, full knowledge of the roles these genes play in normal fish development. The situation is even more confusing because zebrafish lacks an *amhr2* gene (Rocha *et al*. 2016), the loss of which we show here to be associated with chromosomal rearrangements that have breakpoints at the expected site of the ancestral *amhr2* gene, breakpoints that originated at the base of the cypriniform radiation because we show that this inversion breakpoint is shared by the common carp (*Cyprinus carpio*).

To help identify ancestral roles, we knocked out *amh* in the zebrafish *Danio rerio*. We studied gonad development, reproductive tract function, and transcriptomics to help understand the molecular genetic mechanisms of Amh action. Like mammals, zebrafish expresses *amh* in Sertoli cells in testes and in granulosa cells in ovaries (Rodriguez-MARI *et al*. 2005; Von Hofsten *et al*. 2005; Wang and Orban 2007; Chen *et al*. 2017; Yin *et al*. 2017). In adult zebrafish organ culture, Amh inhibited the production of Fsh-stimulated androgen, and also inhibited androgen-stimulated proliferation of spermatogonia (Skaar *et al*. 2011), suggesting a role for Amh in testis function.

Results showed that zebrafish males and females that lack Amh function had enormous gonads due to increased production and/or accumulation of germ cells (Lin *et al*. 2017). Mutant males developed mature sperm able to fertilize eggs, but at lower rates than wild-type siblings. Young mutant females produced fertile eggs, but older females became sterile as their ovaries accumulated immature follicles that failed to deposit yolk. Reproductive ducts in both males and females were structurally and functionally normal, making unlikely the hypothesis that the inhibition of female sex duct development is a conserved feature of Amh across vertebrates. Juvenile *amh* mutant zebrafish developing as males retained oocytes longer than their wild-type siblings, which generally develop as hermaphrodites before transitioning to become males or females about 19 to 30 days post fertilization (dpf) (Takahashi 1977; Rodriguez-MARI *et al*. 2005; Wang *et al*. 2007; Orban *et al*. 2009). This result suggests that Amh promotes oocyte apoptosis in transitioning juvenile zebrafish. Based on trunk transcriptomes, 21dpf transitional stage fish clustered into two groups, one of which expressed more ovary genes, but both groups contained both wild-type and mutant fish, showing that Amh was not playing a sex-specific role at this stage. Transcriptomes of 35dpf juvenile trunks clustered animals into clearly male and female groups, and within each sex group, wild types separated from mutants, showing that at this stage, Amh action is important for gonad development. Transcriptomic comparisons of wild-type and *amh* mutant ovaries and testes revealed an ancestral role of Amh in Leydig cell development, oocyte differentiation, and the regulation of germ cell proliferation. We conclude that Amh either was not important for reproductive duct development in the last common ancestor of zebrafish and humans or, more likely, that this role was lost in the zebrafish lineage along with the loss of Müllerian ducts. A shared role of Amh, however, was likely the inhibition of germ cell proliferation both in ovaries and in testes, and that in mammals, the ovary retained this role but the testis apparently lost it. Alternatively, the teleost lineage gained the male germ cell proliferation role of Amh.

## MATERIALS AND METHODS

### Animals

CRISPR/Cas9 mutagenesis generated deletions in zebrafish *amh* (ENSDARG00000014357, http://ensembl.org) using sites identified by ZiFiT Targeter (http://zifit.partners.org/ZiFiT/). Mutagenesis targeted two regions in *amh* exon-3: GGGATGCTGATAACGAAGGA (Site 1) and GGAATGCTTTGGGAACGTGA (Site 2) using gRNAs synthesized from DNA oligomer templates: aattaatacgactcactataGGGATGCTGATAACGAAGGAgttttagagctagaaatagc and aattaatacgactcactataGGAATGCTTTGGGAACGTGAgttttagagctagaaatagc (IDT, Coralville, IA). MEGAscript T7 Transcription Kit transcribed gRNA and mMESSAGE mMACHINE T3 Transcription Kit (Thermo Fisher Scientific, Waltham, MA) synthesized *Cas9* mRNA. Approximately 2 nl of a solution containing 100 ng/μl *Cas9* mRNA and 25 ng/μl of both *amh* gRNAs was co-microinjected into one-cell embryos of the AB strain. Genomic DNA from injected embryos at 24hpf (hour post-fertilization) provided template to amplify a 319-bp PCR fragment including both sites (primers: `F-AGGGTGTGCATGCTACAGAAGGTAAA and R-TGCCATCTTTTTGCACCATCATTTCCAGCCA`).

Wild-type alleles have an *HpyA*V recognition site at Site 1 and an *HpyCH4*IV recognition site at Site 2 that are disrupted in *amh* mutant alleles. Sanger sequencing (GENEWIZ, Inc. NJ) verified mutations. We established stable lines for three non-complementing alleles: deletions of 5-, 10-, or 26-nucleotides (Fig. 1C, D). In addition, we made TALEN-induced deletions in *amh* (Fig. 1E). TALENs targeted the first coding exons of *amh* and were assembled as previously described (Dranow *et al*. 2016). TALEN RNAs were synthesized by in vitro transcription using the mMESSAGE mMACHINE kit (Ambion). TALEN pairs were co-injected at the one-cell stage at 50-100 pg for each TALEN. Founders were identified by screening sperm DNA by high resolution melt (HRM) analysis (Dahlem TJ *et al*. 2012), using Light Scanner Master Mix (BioFire Defense), a CFX-96 real-time PCR machine and Precision Melt Analysis software (BioRad). Primer sequences for the indicated amplicon used were as follows (wild-type amplicon size in parentheses): F-AGATTTGGGCTGATGCTGAT and R-GTGGGACGAATGACTGACCT (212 bp). After initial identification, subsequent genotyping of offspring was performed by PCR followed by visualization on a 2% agarose gel using the same primers. The mutant allele *amh*(*uc28*) was an 11 bp deletion of the bold-faced nucleotides (<u>ACAGTGAGGCACGAAGA</u>GCAGGACAACAACCCGA<u>AGGTCAACCCGCTATC</u>, with TALEN sequences underlined.

**Figure 1.**
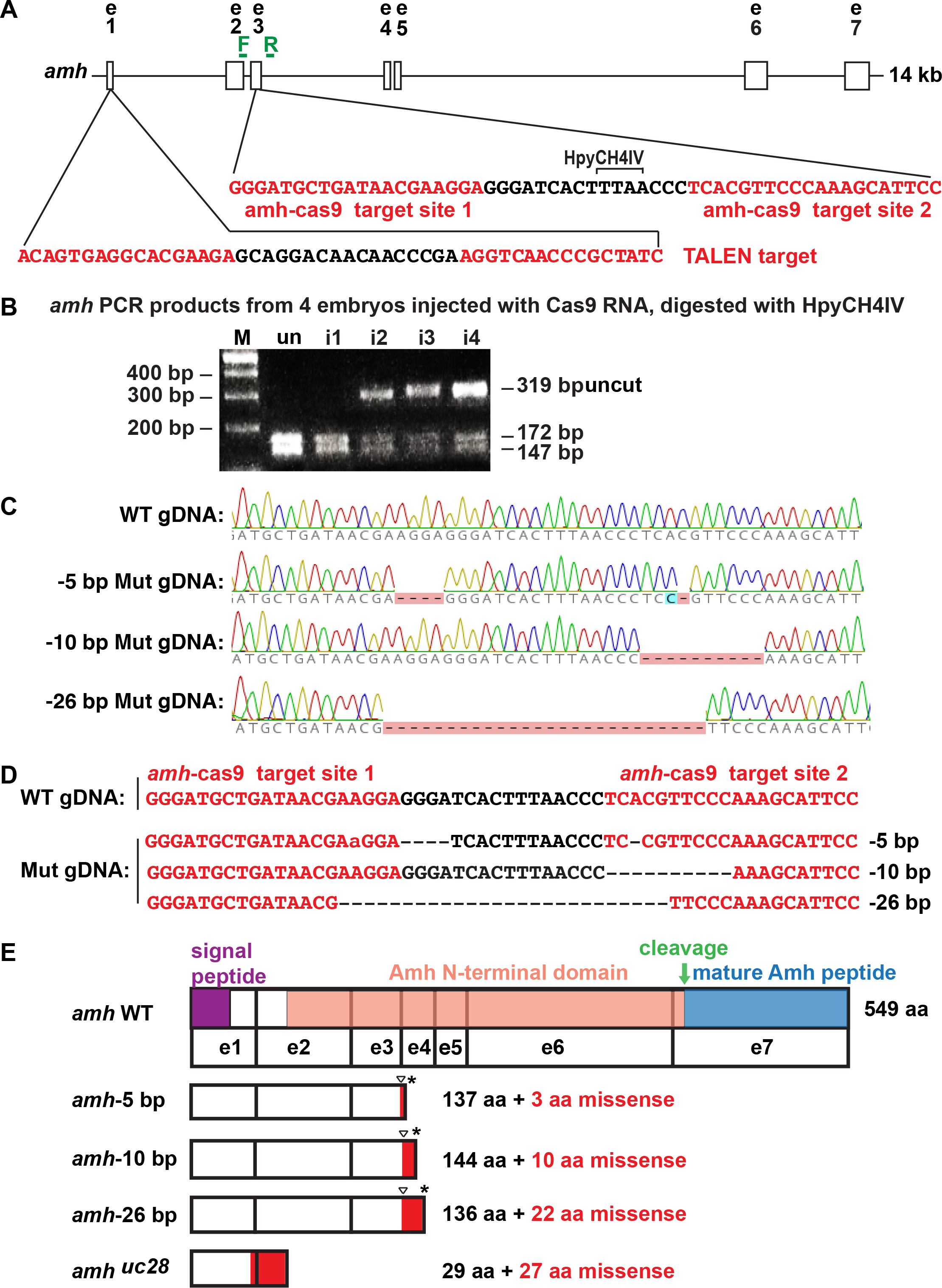
CRISPR/Cas9-induced *amh* mutants. A. 14kb of the *amh* locus showing two CRISPR target sites (red letters) in exon-3. PCR primers, forward (F) and reverse (R) (green). B. Assay for injected CRISPR efficacy. PCR analysis of four G0 injected embryos at 1dpf using genotyping primers F and R shows a 319 base pair (bp) fragment in wild types (WT) that digested with *HpyCH4*IV to produce fragments of 172bp and 147bp; this site disappeared from *amh* genes in a large portion of cells in CRISPR-injected embryos. Abbreviations: M, length marker; un, uninjected 24hpf embryos; i1-i4, CRISPR-injected 24hpf embryos. C. Sequence traces from genomic DNA from a wild-type fish and from three stable mutant lines carrying -5 bp, -10 bp and -26 bp deletions. D. Sequences of genomic DNAs from a wild-type (WT) fish and three stable mutant lines (Mut). E. Predicted structure of Amh protein showing the location of the mutation (triangle), the predicted out-of-frame portion (red), and the premature stop codon (*). Protein coding domains: signal peptide, purple; Amh amino-terminal domain, salmon; cleavage site, green arrow; mature Amh peptide, blue.

### Histology and *in situ* hybridization

*In situ* hybridization was performed as described (Rodriguez-Mari et al., 2005) using the probes: *amh* (ENSDARG00000014357), a 375-bp *amh* fragment including part of exon-7 (primers: F-AGGCTCAGTACCGTTCAGTGTTGC and R-CCAACATCTCCTACAAGACCAACG (Rodriguez-MARI *et al*. 2005)); *bmp15* (ENSDARG00000037491) (Dranow *et al*. 2016); *cyp19a1a* (ENSDARG00000041348) (Chiang *et al*. 2001a); *gata4* (ENSDARG00000098952) using a 763-bp fragment including exon-1 to exon-6 (primers F-AGCACCGGGCACCATCATTCTCCG and R-GAGCTGGAGGATCCGCTTGGAGGC); *gdf9* (ENSDARG00000003229) using a 979-bp fragment including most of the coding region (primers F-TGTTGAACCCGACGTGCCCC and R-TGGTGTGCATTGGCGACCCG); *gsdf* (ENSDARG00000075301) (Yan *et al*. 2017); *bmpr2a* (ENSDARG00000011941) using a 914-bp fragment containing a part of the last coding exon and the 3’UTR (primers bmpr2a + 2,658 F-GAGAGGGAGGAGAGAACAATGAGAGT and bmpr2a -3,572 R-AGGGTACGTATCCACAATAGGTTGGA); *bmpr2b* (ENSDARG00000020057) giving a 727bp fragment is in exons 12 and 13 (primers bmpr2b +2,978 F-GGAGTCTTCGTCGTCTCGATTGAAAT and bmpr2b -3,705 R-TCACCTCTCCGTCTAGTGTATCAGTG); *nr5a1a* (ENSDARG00000103176) using a 859-bp fragment including exon-2 to exon-6 (primers F-AAGTGTCCGGTTATCATTACGGCC and R-TGTCTGCAGATGTGATCCAGAAGC); and *vasa* (ENSDARG00000014373) (Yoon *et al*. 1997). Histology used paraffin-embedded Bouin’s-fixed tissue sectioned at 10 µm and stained with hematoxylin and eosin (H&E) (Rodriguez-MARI *et al*. 2005). GSI, the gonadosomatic index, was calculated as the weight of the gonad divide by the weight of the fish times 100.

### Transcriptomics

Juvenile wild types *and amh^−26^* homozygous mutants at 21 and 35 dpf were euthanized in Tricaine followed by isolating the gonad-containing trunk from just posterior of the pectoral fin to just anterior to the anus. Adult wild-type *and amh^−26^* homozygous mutant gonads were dissected from 8mpf adult animals. Trunks or gonads from each fish were individually homogenized in 200ul Trizol. Total RNA was extracted following (Amores *et al*. 2011), and enriched for mRNA using Dynabeads® Oligo(dt)^25^ (ThermoFisher). We constructed indexed, strand-specific cDNA sequencing libraries (NEXTflex™ qRNA-seq kit, BIOO Scientific), quantified libraries by Qubit® fluorometer (Life Technologies), normalized libraries to 2.3nM, multiplexed and quality checked libraries (Kapa Library Quantification Kit, Kapa Biosystems), and sequenced them in one lane on an Illumina HiSeq 4000 (paired-end 100 base pairs (bp)).

### Bioinformatics

The Dupligänger duplicate removal pipeline (Sydes *et al*. 2019) preprocessed RNA-seq reads, identified and removed BIOO inline UMIs from the 5ʹ-end of each read, removed read-through adapters (cutadapt v1.15 (Martin 2011), command line options: -n 3 -O 1 -m 30 -a AGATCGGAAGAGC -A AGATCGGAAGAGC --too-short-output --too-short-paired-output), and then removed low quality sections from both the 5ʹ-ends and 3ʹ-ends (Trimmomatic (v0.36) (Bolger *et al*. 2014), command line options: LEADING:10 TRAILING:10 SLIDINGWINDOW:5:10 MINLEN:30).

Dupligänger tracked the number of nucleotides removed from the 5ʹ-end and removed reads shorter than 30nt. We aligned processed PE reads to the zebrafish genome (GRCz10, Ensembl version 91) in a splice-aware manner using GSNAP (Wu *et al*. 2016) (v2017-06-20, command line options: --suboptimal-levels 0 --quiet-if-excessive --kmer 15 --max-mismatches 0.1 --use-splicing -- split-output), retaining reads that aligned in a concordant and unique manner. Dupligänger then removed PCR duplicates from the sequence alignment file if both of the following criteria had already been observed in another read pair: a) the read pair shares 5’ alignment starts for both R1 and R2 after correcting for 5’ trimming, and b) the read pair shares the same R1 UMI and R2 UMI. We passed de-duplicated sequence alignment files to HTSeq-count (Anders *et al*. 2015) (command line options: --mode intersection-strict --type exon --stranded reverse) to obtain per-gene counts for protein-coding genes. DESeq2 provided statistical analysis of fold changes (Love *et al*. 2015). Analysis of conserved syntenies used the Synteny Database and Genomicus (Catchen *et al*. 2009; Nguyen *et al*. 2018).

### **D**ata availability statement

RNA-seq reads are available at the Sequence Read Archive (https://www.ncbi.nlm.nih.gov/sra) under accession number PRJNA512103. Supplemental TableS1 (TableS1 Transcriptomics amh Juvenile trunks 21&35dpf.xls) and Supplemental Table S2 (TableS2 Transcriptomics amh Adult gonads.xls) list differentially expressed genes for juvenile trunks or adult gonads for *amh* mutants and wild-type siblings, respectively. Work was performed under the University of Oregon IACUC protocol #14-08R. Mutant strains are available on request.

## RESULTS

### Molecular genetics of induced *amh* mutations

To identify the roles of Amh in gonad development, we induced frame-shift premature stop codon alleles in zebrafish *amh* (ENSDARG00000014357) using CRISPR/Cas9 and TALEN mutagenesis. CRISPR guide RNAs targeted two sites in exon-3 located 16 nucleotides (nt) apart (Fig. 1A, red). These sites should be translated into the protein’s Amh-domain, upstream of the cleavage site that liberates the TGF-beta domain that encodes the mature functional Amh protein. To assay CRISPR efficacy, we injected gRNAs and Cas9 RNA into 1-cell AB strain embryos, and at 24hpf (hours post fertilization), extracted DNA, amplified the target (primer locations green in Fig. 1A), and digested fragments with *HpyCH4*IV, which cleaves the wild-type but not a mutated site. Three of the four embryos tested had substantially reduced restriction enzyme cleavage (Fig. 1B), verifying reagent utility. We raised injected embryos and isolated three mutant lines. Sanger sequencing (Fig. 1C) revealed deletions of 5, 10, and 26 nt (Fig. 1D, designated below as CRISPR-induced alleles *amh^−5^*, *amh^−10^*, and *amh^−26^*) and a deletion of 11 nt as a TALEN-induced *amh*(*uc28*) allele. These frame-shift mutations should result in truncated proteins lacking the mature TGF-beta domain due to premature stop codons (Fig. 1E).

### Amh facilitates development of a male phenotype

To learn if *amh* plays a role in zebrafish sex determination as in some other fish (Hattori *et al*. 2012; Kamiya *et al*. 2012; Li *et al*. 2015), we investigated sex ratios in *amh* mutant lines. The sex ratio of wild-type siblings was unbiased (48.4% males, 41 males and 43 females), but homozygous mutants had an average of only 17.8% males (12 males and 54 females, p < 0.05, Wilcoxon Rank Sum Test), about a third as many as expected, similar to prior results (Lin *et al*. 2017). We conclude that wild-type *amh* functions to facilitate the development of males but is not essential for AB strain zebrafish to develop a male phenotype.

### Amh regulates the production of functional gametes

To test female fertility, we mated individual *amh* mutant females (−26 allele) to AB wild-type males and to test male fertility, we mated individual *amh* mutant males (−26 allele) to AB wild-type females. For both tests, we counted the number of females that laid eggs, the number of eggs per clutch, and the number of embryos that developed up to 72hpf. Results showed that homozygous *amh* mutant females at 4.5 months post fertilization (mpf) laid about half as many eggs as wild types (87±57 eggs/cross vs. 169±100 eggs/cross), but most eggs from mutant females supported normal embryonic development (744/961 eggs, Fig. 2A). Homozygous *amh* mutant females at 11mpf failed to lay any eggs at all (Fig. 2B). These results show that although young *amh* mutant females laid fewer eggs than normal, they nevertheless did lay eggs that developed; we conclude that *amh* mutant females developed functional reproductive ducts and results suggest that Amh is necessary for continued fertility as zebrafish age.

**Figure 2.**
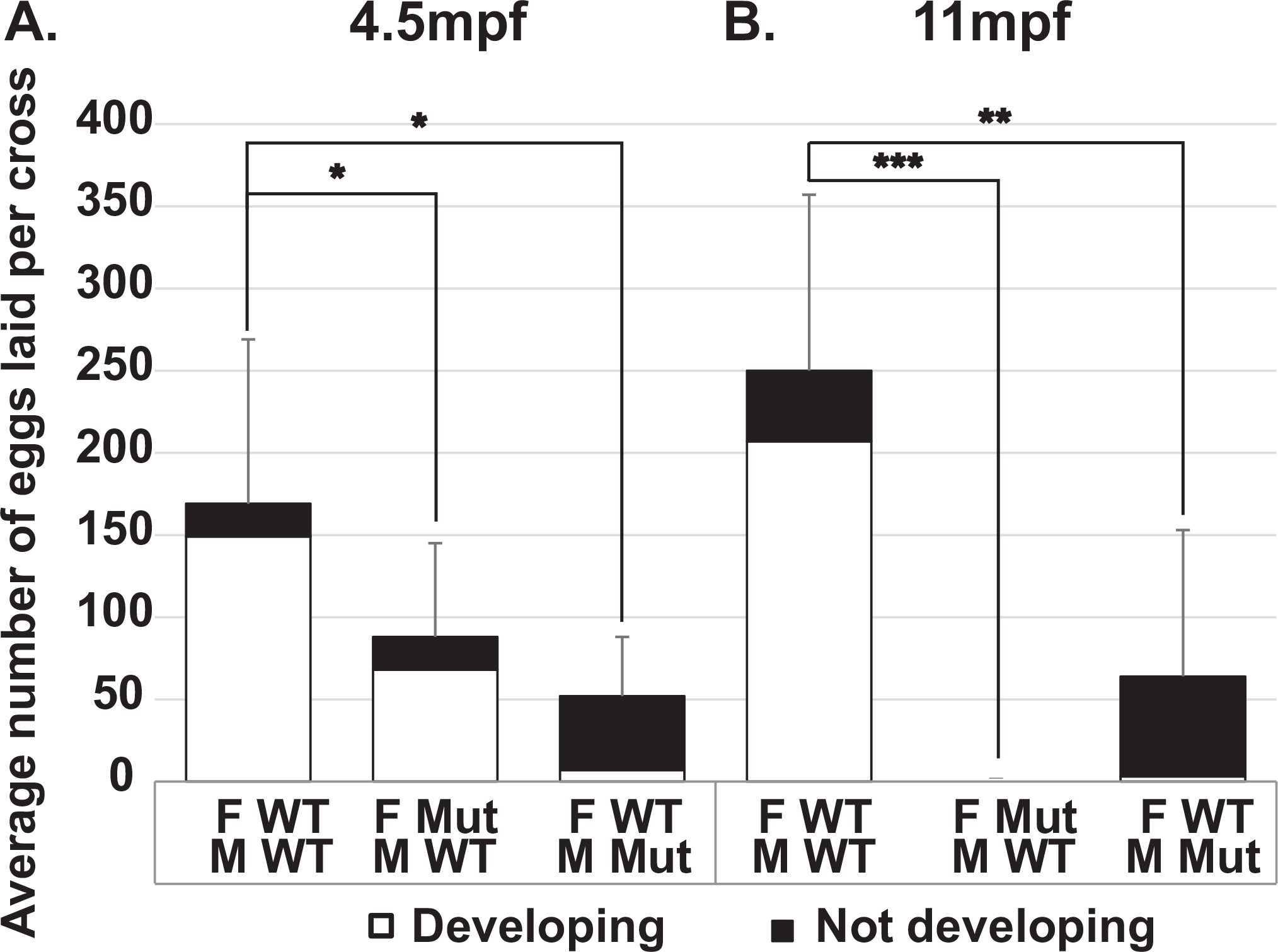
Fertility tests for adult *amh* mutants and wild types. (A) Average number of eggs laid per cross from wild-type females crossed to wild-type males (11 crosses), *amh^−26^* mutant females crossed to wild-type males (11 crosses), and wild-type females crossed to *amh^−26^* mutant males (4 crosses) at 4.5mpf. (B) Average number of eggs laid per cross from wild-type females crossed to wild-type males (8 crosses), *amh^−26^* mutant females crossed to wild-type males (7 crosses), and wild-type females crossed to *amh^−26^* mutant males (6 crosses) at 11mpf. For each cross, one individual female (either mutant or wild-type sibling) was paired with three non-sibling wild-type males, or for the reciprocal test, one individual male (either mutant or wild-type sibling) was paired with three non-sibling wild-type females. Eggs were collected and counted at 1dpf and 3dpf; embryos were scored as developing normally (white bars), or as not developing or improperly developing (black bars). Statistical significance: *, 0.05<p<0.01; **, 0.01<p<0.001 and ***, p< 0.001, Wilcoxon Rank Sum Test. Error bars: standard deviation. Abbreviations: F, female; M, male; WT, wild type; Mut, mutant.

Tests of *amh* mutant male fertility showed that at 4.5mpf, crosses of single *amh*^−26^ homozygous mutant males by three wild-type females resulted in the laying of only about 27% as many eggs as did wild-type sibling males (45±36 eggs/cross vs. 169±100 eggs/cross, respectively), suggesting that normal Amh activity improves male mating behaviors. Only about 11% of eggs (5±4 of 45±36) laid by wild-type females mated to mutant males initiated development (Fig. 2A), showing that Amh is required for optimal sperm production and/or function. Results for homozygous *amh^−26^* mutant males at 11mpf showed continuing severe effects on male fertility (Fig. 2B). These results indicate that young mutant males make and release mature functional sperm, and thus that their reproductive ducts can transport sperm, at least initially. We conclude that *amh* function is not required for normal male sex duct development but is necessary for normal rates of functional sperm production. Combined with results from mutant females, we conclude that Amh is not required to construct functional reproductive ducts or to initiate fertility in either sex but is necessary to maintain fertility in both sexes.

### Amh promotes juvenile gonad development

To understand *amh* mutant gonadal phenotypes, we studied histological sections at several developmental stages. For 21dpf late larval zebrafish, all eight wild-types examined had gonads with stage I oocytes (Selman *et al*. 1993; Maack 2003) as expected for zebrafish juvenile hermaphrodites (Takahashi 1977; Rodriguez-MARI *et al*. 2005; Wang and Orban 2007; Rodriguez-MARI *et al*. 2010). Fig. 3A and B show two of the eight individuals. Six of eight 21dpf *amh^−26^* mutants were similar to wild types with stage I oocytes (Fig. 3C), but two lacked stage I oocytes and contained only undifferentiated germ cells (Fig. 3D). We conclude that most *amh* mutants develop histologically normal gonads at 21dpf, although some have gonads with delayed development.

**Figure 3.**
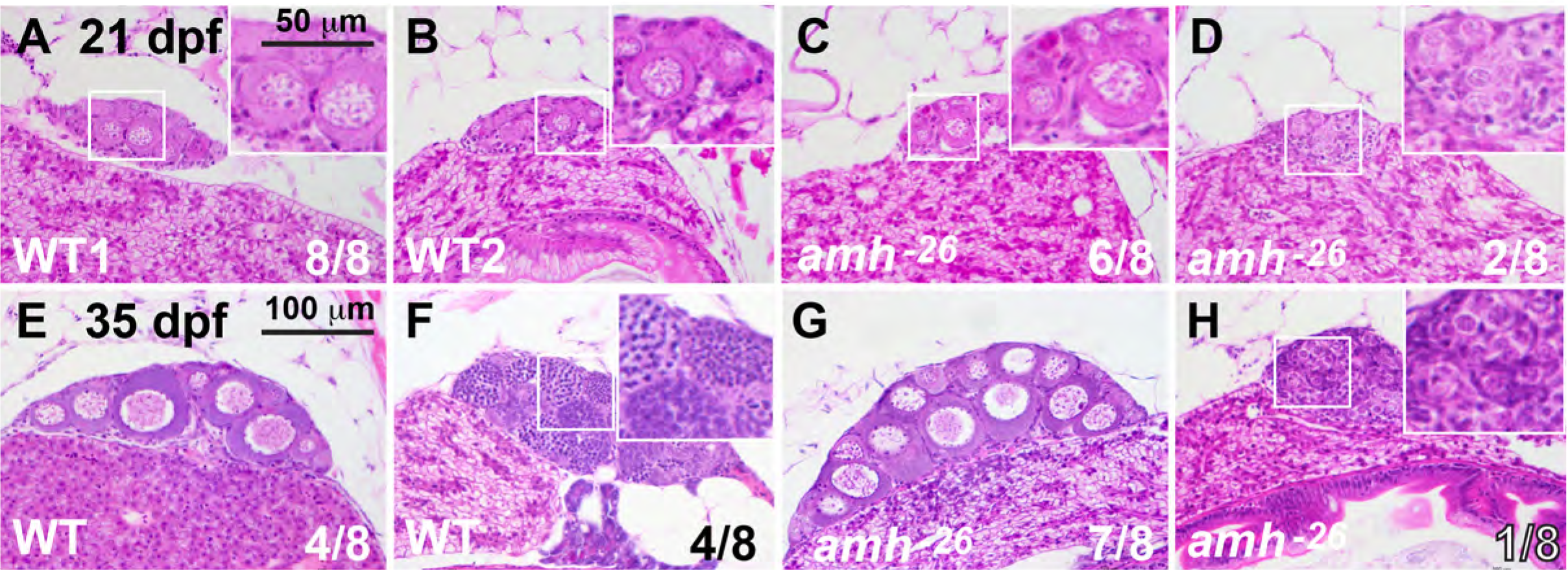
Gonad histology of 21dpf and 35dpf wild-type and *amh-*mutant fish. A-D. In histological sections, gonads in all eight 21dpf wild-type sibling fish (WT) contained early oocytes (one gonad shown in each of two individuals in A and B). Gonads in six of eight 21dpf *amh* ^−26^ mutants were morphologically like wild-type ovaries (C) and gonads of two of eight 21dpf *amh* ^−26^ mutants were undifferentiated (D). At 35dpf, wild-type fish contained gonads that were clearly either ovaries (4/8 fish) (E) or testis (4/8 fish) (F). In 35dpf *amh* ^−26^ mutants, most fish had ovaries (7/8 fish) (G) but one of eight fish had immature testis (H). Smaller boxed regions in several panels are magnified in the larger boxed regions at the right of these panels. Scale bar in E is 100µm for all panels; scale bar in the higher magnification boxes in A: 50 µm.

For 35dpf juveniles, four of the eight wild types examined had stage I-II oocytes (Fig. 3E) and four had developing spermatocytes and spermatozoa (Fig. 3F). Among the eight *amh* mutants examined, seven had ovaries with morphologies similar to those in wild types (Fig. 3G) and only one fish had gonads that lacked oocytes and possessed developing spermatogonia organized in cysts (Fig. 3H) (Maack 2003). We conclude that most of the 35dpf mutant juveniles we examined were embarking on a female trajectory, and that the only 35dpf *amh* mutant male that we sectioned had gonads that were developmentally delayed with respect to those in wild-type siblings.

### In females, Amh inhibits germ cell proliferation and differentiation

To learn the roles of Amh in adults, we investigated gonad morphology in *amh* mutants over time. In adult females at 8mpf, ovaries in wild-type siblings contained oocytes of all stages (Fig. 4A, E, I). In contrast, *amh^−26^* mutant females had enlarged ovaries that distended the individual’s abdomen (Fig. 4B, F, J). Averaging results from females homozygous for the *amh*^−10^ and *amh*^−26^ alleles, the gonadal-somatic index (GSI, (gonad weight / body weight)*100) of *amh* mutants was about 2.6-fold larger than their respective wild-type siblings, confirming prior results in zebrafish and mouse (Durlinger *et al*. 1999; Lin *et al*. 2017). Adult ovaries in 8mpf *amh^−^*^26^ zebrafish mutants lacked oocytes that had matured beyond stage III (Fig. 4F, J). Young (4.5mpf) *amh^−26^* mutant ovaries had 2.7-times as many stage I and II oocytes as found in wild-type ovaries (Fig. 4W), and by 8mpf and 18mpf, the relative proportion of immature oocytes increased to 9- and 35-fold that in wild-type siblings, respectively (Fig. 4W). We conclude that Amh activity inhibits oogonia proliferation or maturation. Although young *amh^−^*^26^ mutants had formed stage IV oocytes in the central gonad (average of 23 stage IV oocytes in mutants and 32 in wild types), 8mpf *amh^−^*^26^ mutant females had few stage IV oocytes in the central gonad (average of 5 oocytes in mutants and 20 in wild types) and 18mpf *amh-*^26^ mutant females had an average of only two stage IV oocytes vs. 17 in wild types (Fig. 4W). Homozygotes for the *amh*^−5^, *amh*^−10^, and *amh^uc28^* alleles displayed similar phenotypes (Supplemental Fig. S3). We conclude that in aging female zebrafish, Amh activity is required to advance ovarian follicles from stage III to more mature stages.

**Figure 4.**
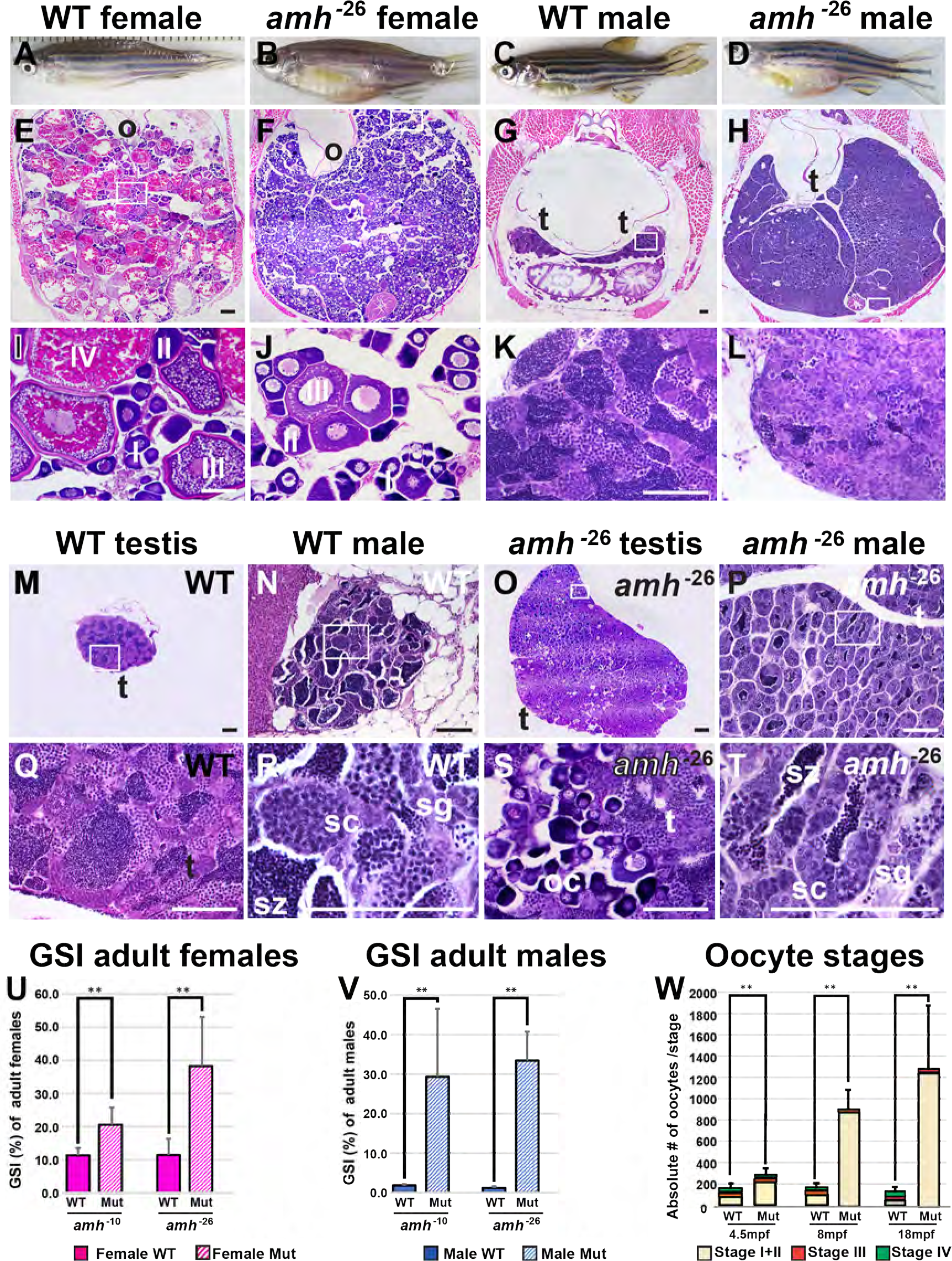
Amh activity is required for normal gonad morphology in adult zebrafish. (A-D) 8mpf adult zebrafish: wild types (A, female, 6 fish sectioned; C, male, 7 fish) and *amh^−26^* mutants (B, female, 6 fish; D, male, 7 fish), showing enlarged abdomens in mutants. (E-T) Histological sections of 8mpf adult gonads: Adult female ovaries at low (E, F), and high (I, J) magnification. Cross-sections of an 8mpf wild-type female sibling (E, I) revealed maturing (stage-I and -II) and vitellogenic (stage-III and -IV) follicles. Cross-sections of an 8mpf *amh* mutant female (F, J) showed an excess of immature follicles (stage-I and -II), a few early vitellogenic follicles (stage-III), but no late vitellogenic follicles (stage-IV) (Numbers of oocytes per stage shown in panel W). Panels M-T illustrate some of the variation in mutant phenotypes, (M, Q) Low and high magnification of dissected wild-type testis. (N, R) Medium and high magnification of a cross section of the abdomen of a different wild-type male. (O, S) Low and high magnification of dissected ovotestis from an *amh^−26^* mutant male showing immature oocytes in the testis. (P, T) Medium and high magnification of the abdomen of a different *amh^−26^* mutant male showing small testis lobules and fewer late stage male gonocytes compared to wild types. Gonadal somatic index (GSI) of adult females (U) and males (V). GSI calculations for females used five wild-type siblings of *amh^−10^* females, five *amh^−10^* mutant females, five wild-type siblings of *amh^−26^* mutant females, and five *amh^−26^* mutant females. GSI calculations for females used five wild-type siblings of *amh^−10^* males, five *amh^−10^* mutant males, eight wild-type siblings of *amh^−26^* mutant males, and five *amh^−26^* mutant males. (W) Number of oocytes per stage at 4.5, 8, and 18mpf. Oocytes were categorized into three groups: stage I + stage II (beige), stage III (red), and stage IV (green) oocytes in W. The 18mpf mutant females had mostly stage I + stage II oocytes (W). Statistical significance: **, 0.01<p<0.001 and ***, p< 0.001; Wilcoxon Rank Sum Test). In (U) and (W), solid boxes, wild-types; striped boxes, mutants; red boxes, females; blue boxes, males. Abbreviations: I, II, III, IV: ovarian follicle stages 1 to 4; o, ovary; s, Sertoli cells; sc, spermatocytes; sg, spermatogonia; sz, spermatozoa; t, testis. Black scale bar in E for E and F; black scale bar in G for G and H; white scale bar in I for I and J; white scale bar in K for K and L. All scale bars: 100µm. (U, V and W) Gonadosomatic index (GSI) in percent. (U): wild types (WT) and *amh* mutant (Mut) ovary. (V) wild types (WT) and *amh* mutant (Mut) testis.

### In males, Amh inhibits germ cell proliferation and oocyte development or survival

Males homozygous for each of the four *amh* mutant alleles displayed several phenotypic differences from wild-type siblings at 8mpf. First, *amh* mutant males had much larger abdomens than wild-type siblings (Fig. 4C, D) due to greatly enlarged testes (Fig. 4G, H, K. L), confirming prior results (Lin *et al*. 2017). The overgrowth of *amh* mutant male gonads (about 33.7 times heavier than wild-type sibling gonads (an average of 0.207±0.103 g (SD) for mutant testes (n=10) vs. 0.006±0.003 g for wild-type testes (n=13)) was even larger than that of mutant female gonads (2.2 fold, an average of 0.171±0.078 g for mutant ovaries (n=10) vs. 0.078±0.027 g for wild-type ovaries (n=10)) (see Fig. 4V). We conclude that *amh* activity is required to inhibit gonad growth both in adult males and in adult females. Adult *amh* mutant male gonads contained all stages of sperm development, including mature spermatozoa (Fig. 4L, P, T). Second, the proportion of later stage male gametocytes in *amh* mutant testes appeared to be greatly reduced compared to wild types and the proportion of immature stages seemed much higher in mutant males than wild type males (Fig. 4L vs. Fig. 4K). Third, testis tubules were smaller in size but greater in number in *amh* mutants compared to wild types (Fig. 4 G, H, K, L, M-T). In cross sections, lobules in mutant testes were only 19.3% as large as lobules in wild-type testes (638± 272 vs. 3,313± 611 µm^2^). Fourth, and most remarkable, more than half of the 8mpf *amh^−26^* male mutant gonads examined (4/7 fish) contained early stage oocytes, but none of the seven 8mpf wild-type male siblings did (Fig. 4M-T). The finding of ovo-testes in mature adult *amh* mutants shows that normal *amh* activity helps to masculinize zebrafish gonad development by inhibiting the production or survival of young oocytes. We conclude that in zebrafish, normal Amh activity is required to regulate the proliferation of spermatogonia, to control the number and size of testis tubules, to govern the rate of maturation of spermatogonia to spermatozoa, and to ensure that immature oocytes disappear from male gonads during the juvenile hermaphrodite stage or to block the formation of oocytes in later development.

### Amh and Gsdf appear to act in the same developmental pathway

Gsdf, like Amh, is essential to prevent the accumulation of young oocytes as zebrafish females age (Yan *et al*. 2017). If these two genes act in the same pathway, then double mutant ovaries should have about the same phenotype as each single mutant. Alternatively, if the genes act in parallel pathways, then double mutants should have more severe phenotypes than either single mutant. Analysis of *amh;gsdf* double mutants revealed female gonad phenotypes that were about the same as in each of the two single mutants: all three genotypes accumulated an enormous number of small oocytes with few stage III oocytes at 8-12mpf (Supplemental Fig. S1 and data not shown). Males homozygous mutant for either *amh* or *gsdf* had enlarged testes compared to wild types; *amh* mutant males had larger testes even than *gsdf* mutant males; and *amh* mutant males became sterile as they aged while *gsdf* mutant males maintained fertility. Double mutant testes were similar to *amh* mutants, and not more severe (Supplemental Fig. S1), consistent with the explanation that in males as in females, *amh* and *gsdf* act in the same pathway.

Furthermore, *amh e*xpression was nearly twice as high in *gsdf* mutant testes as in wild type testes (Yan *et al*. 2017), suggesting that Gsdf controls *amh*. Reciprocally, *gsdf* expression was 3.4-fold higher in *amh* mutant adult testes compared to wild-type testes in our RNA-seq results (see Suppl Table 2), consistent with the result from *in situ* hybridization (Fig. 5H and 5H’), suggesting that Amh controls *gsdf*. Together the mutant phenotypes and expression data show that the regulation of these two TGFb family genes are interdependent.

**Figure 5.**
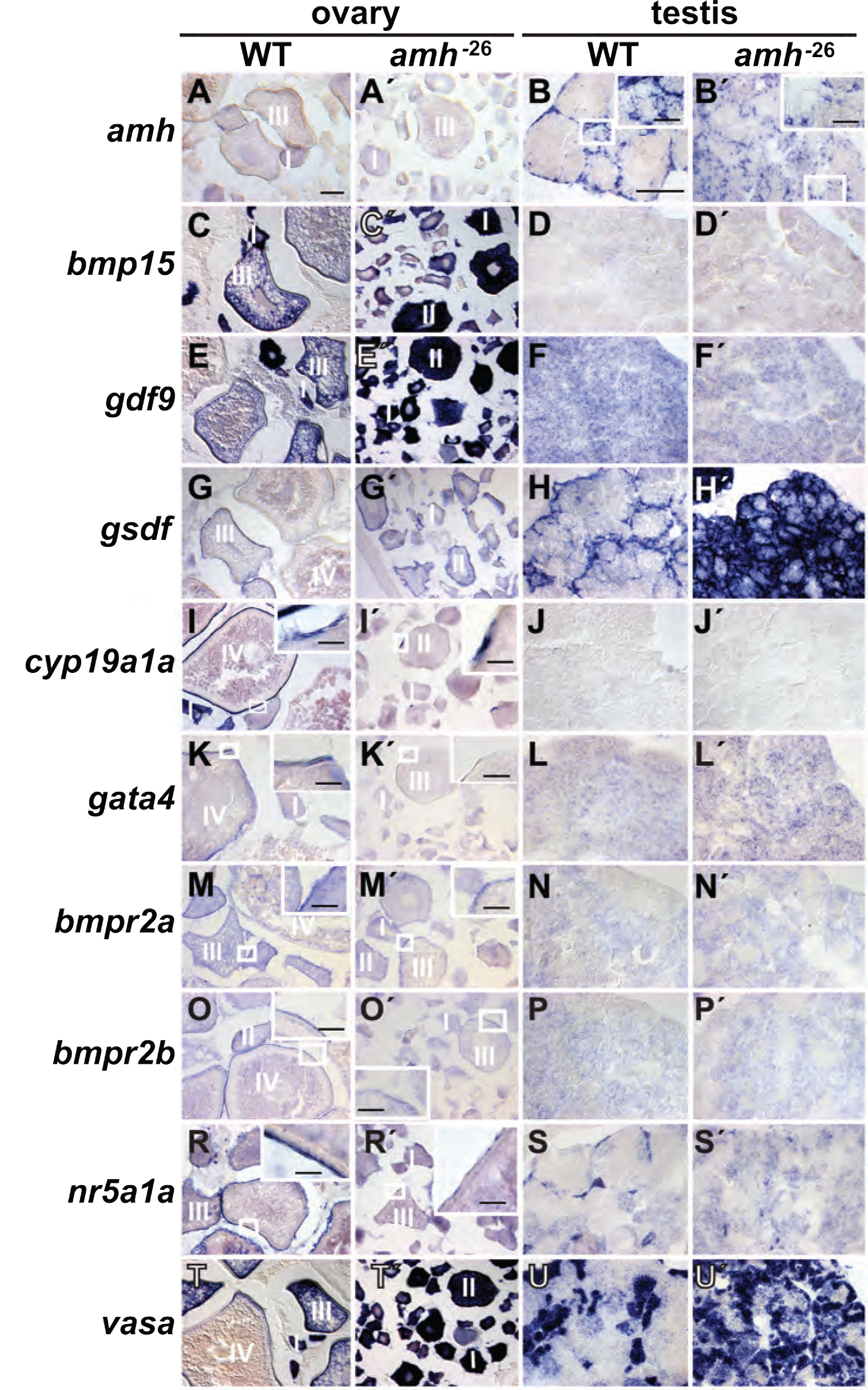
Gene expression patterns in adult gonads at 8mpf. Wild-type ovaries (A, C, E, G, I, K, M, O, R, T); *amh*^−26^ mutant ovaries (Á, Ć, E′, G′, Í, K′, M′, Ó, R’ T’); wild-type testis (B, D, F, H, J, L, N, P, S, U); *amh* mutant testis (B′, D′, F′, H′, J′, Ĺ, N′, P′, S’, U’). *In situ* hybridization for *amh* (A, Á, B, B′), *bmp15* (C, Ć, D, D′), *gdf9* (E, E′, F, F′), *gsdf* (G, G′, H, H′), *cyp19a1a* (I, Í, J, J′), *gata4* (K, K′, L, Ĺ), *bmpr2a* (M, M’, N, N’), *bmpr2b* (O, O’, P, P’), *nr5a1a* (R, R’, S, S’), *vasa* (T, T’, U, U’). Small boxed regions in low magnification views are shown in larger boxed regions at higher magnification for B, B′, E, E′, G, I, I’ K, K’, M. M’, O, O’, R, R’ M, M′. Scale bars for main panels represent 100µm; scale bars for higher magnification in boxed regions represent 25µm. Abbreviations: I, II, III, IV: ovarian follicle stages 1 to 4.

### Amh activity is required for normal expression of key gonad development genes

To understand in more detail the role of Amh in zebrafish gonad development, we studied the expression of several key regulatory and marker genes in adult wild types and *amh* mutants by *in situ* hybridization. Wild-type adult ovaries at 8mpf expressed *amh* mainly in granulosa cells surrounding stage II oocytes (Fig. 5A, see also (Rodriguez-MARI *et al*. 2005; Von Hofsten *et al*. 2005)). In contrast, *amh*^−26^ mutant ovaries at 8mpf showed little *amh* expression in somatic cells surrounding oocytes, due either to nonsense-mediated decay or to the failure of *amh-*expressing cells to form in *amh* mutants (Fig. 5A’). Wild-type males at 8mpf displayed a well-organized pattern of *amh* expression in Sertoli cells surrounding testis tubules (Fig. 5B, see also (Rodriguez-MARI *et al*. 2005; Von Hofsten *et al*. 2005)). Presumptive Sertoli cells also expressed *amh* in 8mpf *amh* mutant males, demonstrating transcript stability, but *amh-*expressing cells were less organized; testis tubules appeared to be smaller; and *amh*-expressing cells did not completely surround most testis tubules (Fig. 5B, B’). Homozygous *amh*^−5^ and *amh*^−10^ mutants showed similar expression patterns (data not shown). We conclude that in adult male zebrafish, *amh* is required for the organization of Sertoli cells in testis tubules.

Bmp15 is an extracellular signaling protein that 8mpf wild-type adult zebrafish express mainly in oocytes in early stage ovarian follicles, and in maturing oocytes in later stage wild-type follicles (Fig. 5C and (Clelland *et al*. 2006; Dranow *et al*. 2016)). Adult *amh* mutant ovaries appeared to express *bmp15* stronger than wild-type ovaries did (Fig. 5C, Ć) due to the accumulation of younger stages that express high levels of *bmp15*. Neither wild-type nor mutant testes showed significant *bmp15* expression (Fig. 5D, D′). We conclude that Amh function promotes the maturation of ovarian follicles in mature adult ovaries.

Gdf9, like Bmp15, is a TGF-beta-family member that marks oocytes (Liu and Ge 2007; Dranow *et al*. 2016). Expression of *gdf9* appeared to increase in mature adult *amh* mutant ovaries compared to wild-type ovaries (Fig. 5E, E′), likely due to accumulating young oocytes in *amh* mutants. Testes showed negligible *gdf9* expression in either *amh* mutants or in wild-type siblings (Fig. 5F, F′). We conclude that *amh* function is necessary for the maturation of oocytes to stages in which they appear to down-regulate *gdf9* transcript.

Gsdf is an important signaling molecule in fish gonadogenesis (Rondeau *et al*. 2013; Imai *et al*. 2015; Zhang *et al*. 2016). Wild-type ovaries express *gsdf* in granulosa cells surrounding oocytes (Fig. 5G, (Gautier *et al*. 2011a; Yan *et al*. 2017)). Zebrafish *amh* mutant ovaries also expressed *gsdf* in epithelial cells surrounding immature oocytes (Fig. 5G′). Testes expressed *gsdf* specifically in Sertoli cells surrounding germ cells (Fig. 5H, (Gautier *et al*. 2011a; Yan *et al*. 2017)). Testes lacking *amh* activity showed substantially greater *gsdf* expression than normal, suggesting altered Sertoli cell development (Fig. 5H, H′). Expression of the Sertoli cell marker *amh* in *amh* mutant testes showed that Sertoli cells were poorly organized with smaller testis tubules (Fig. 5B, B’), which was confirmed by *gsdf* expression (Fig. 5H, H’) and histology (Fig. 4N, P). Taken together, results from *gsdf* expression and histology analyses show that *amh* mutants appeared to have many more testis tubules, but much smaller testis tubules, than normal, consistent with an increase in Sertoli cells or their precursors. We conclude that Amh function in adult male zebrafish is necessary for the organization and number of *gsdf*-expressing cells and may help regulate *gsdf* expression.

Aromatase, encoded in zebrafish ovaries by *cyp19a1a* (and in the brain by *cyp19a1b* (Chiang *et al*. 2001a; Chiang *et al*. 2001b)), converts testosterone to estrogen (Rouiller-FABRE *et al*. 1998). As in humans, adult wild-type zebrafish express *cyp19a1* in granulosa cells and theca cells in ovarian follicles (Fig. 5I, (Chiang *et al*. 2001a; Chiang *et al*. 2001b; Dranow *et al*. 2016)). In contrast, young stage follicles in adult zebrafish *amh* mutant ovaries showed fewer *cyp19a1a* expressing cells in patches that did not completely surround follicles (Fig. 5I′). In testes, *cyp19a1a* expression was not detected in either wild types or *amh* mutants (Fig. J, J′). We conclude that *amh* activity is required for ovarian follicles to advance to the strongly aromatase-expressing stage and for the organization of granulosa cells around ovarian follicles.

GATA4 in human gonads synergistically activates the *AMH* promoter by interacting with NR5A1 (SF-1), a process necessary for normal human sex development (Lourenco *et al*. 2011). In mouse, granulosa cells and theca cells express *Gata4* (Padua 2014) and in wild-type zebrafish, oocytes express *gata4* in early stages and granulosa and theca cells express *gata4* in later stages (Fig. 5K, (Yan *et al*. 2017)). In zebrafish, adult female *amh* mutants, like wild-types, dispayed *gata4* transcript in young oocytes, but it was patchy in follicular cells due presumably to alterations in follicular maturation (Fig. 5K′). Expression of *gata4* was low in both wild-type and mutant adult testes (Fig. 5L, Ĺ). These results suggest that *amh* activity normally helps to up-regulate *gata4* in granulosa cells of wild-type ovaries.

Bmpr2 is likely the type II receptor for BMP15 (Moore *et al*. 2003; Pulkki *et al*. 2012). Zebrafish has two co-orthologs of *Bmpr2*: *bmpr2a* is expressed in young oocytes and ovarian follicle cells and *bmpr2b* is expressed in follicle cells (Li and Ge 2011; Dranow *et al*. 2016). Our *in situ* hybridization experiments confirmed the wild-type expression pattern of *bmpr2a* and showed that in mutant ovaries, *bmpr2a* expression was reduced in young oocytes but was maintained weakly in stage III follicles (Fig. 5M, M’). For *bmpr2b,* expression appeared in wild types in follicle cells, but in *amh* mutants, reduced signal was detected in follicle cells (Fig. 5O, O’). Testes in both wild types and *amh* mutants appeared to possess little expression of either *bmpr2* gene and no difference appeared to distinguish wild types from mutants (Fig. 5 N, N’, P, P’).

NR5A1 (alias steroidogenic factor 1, SF-1) interacts with Gata4 protein in cultured primary rat Sertoli cells to up-regulate *Amh* expression (Tremblay 2001). Zebrafish adult ovaries express *nr5a1a* (Von Hofsten *et al*. 2005), and our *in situ* studies showed that this expression is in granulosa cells (Fig. 5M), as it is in mammals. Adult *amh* mutant females expressed *nr5a1a* in a much-reduced and fragmented, patchy, granulosa cell layer (Fig. 5M′), showing that Amh is important for the organization or development of granulosa cells. Adult wild-type testes expressed *nr5a1a* in Leydig cells (Fig. 5N), but far fewer cells expressed *nr5a1a* in mutant testes compared to wild-type testes (Fig. 5N′), despite the increased number of testis tubules in *amh* mutants (compare *nr5a1a* expression in Fig. 5R, Ŕ, to *gsdf* expression in Fig. 5H, H′). We conclude that in male zebrafish, *amh* function is required for normal Leydig cell development. These results show that in both male and female adult zebrafish, cells expressing *nr5a1a* require *amh* function for normal development, and, because Nr5a1 and Gata4 proteins interact to control *Amh* expression in mammals (Tremblay 2001; Lourenco *et al*. 2011), these three genes likely act in a feedback loop.

Vasa, a putative RNA helicase encoded by *ddx4,* is expressed in germ cells in wild-type zebrafish (Fig. 5T, U, (Yoon *et al*. 1997)). Zebrafish *amh* mutants also expressed *ddx4* in germ cells in both males and females (Fig. 5T’ and U’). The intensity of *vasa* signal in wild-type oocytes diminished as follicles matured (Fig. 5T, (Yoon *et al*. 1997)), but in adult *amh* mutants, all oocytes showed high levels of *vasa* expression, consistent with a failure of oocyte maturation in *amh* mutant ovaries (Fig. 5T, T’). In adult testes, *amh* mutants appeared to have more, but smaller, groups of germ cells than did wild-types (Fig. 5U, U’). We conclude that differences in *ddx4* expression reflect the histological differences between wild-type and *amh* mutant gonads.

### Zebrafish *amh* mutants help to identify gene regulatory pathways in gonad development

To help understand genetic programs that regulate gonad development, we sequenced 45 strand-specific RNA-seq libraries, each sample derived from a single individual fish at one of three different ages. 1) Fifteen samples comprised the gonad-containing trunks of 21dpf transitional state juveniles (eight wild types and seven *amh*^−26^ mutants). 2) Another 15 trunks were from 35dpf juveniles (eight wild types and seven *amh*^−26^ mutants. 3) The final 15 libraries came from mature adults at 8mpf, including seven pairs of testes (three individual wild types and four different *amh* mutants) and eight pairs of ovaries (four wild types and four *amh*^−26^ mutants). These 45 RNA-seq libraries produced 396 million paired-end sequence reads, of which 211 million mapped to the Ensembl v91 protein-coding exons of the zebrafish GRCz10 version of the zebrafish reference genome. Two-way similarity clustering (rlog (regularized log) transformed Euclidean distances)) of all samples produced a clear separation between young juveniles, older juveniles, adult ovaries, and adult testes (Fig. 6).

**Figure 6.**
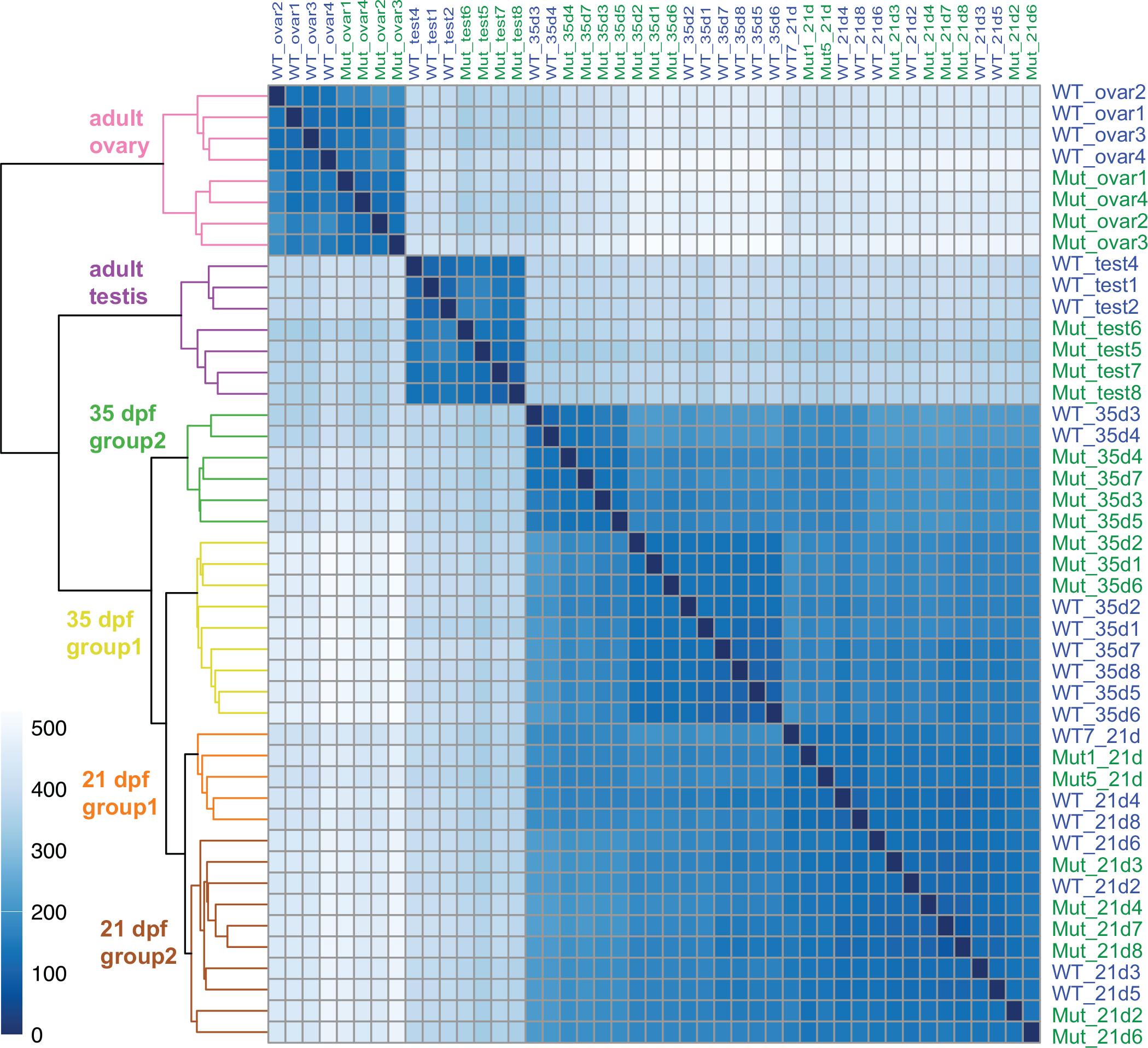
Heat map and dendrogram of rlog-transformed Euclidean distances between all 45 RNA-seq samples. Analysis divided samples into six groups: adult ovary, adult testes, two groups of 35dpf trunks, and two groups of 21dpf trunks. The intensity of each cell in the panel reflects the number of genes different in the intersecting two samples according to the scale at the left, so the diagonal self-comparisons show no genes differently expressed.

### Genome-wide transcriptomics of wild-type adult zebrafish ovaries

Interpretation of gene expression changes in developing mutant gonads requires knowledge of gene expression patterns in adult wild-type gonads (Santos *et al*. 2007a; Santos *et al*. 2007b; Sreenivasan *et al*. 2014; Lee *et al*. 2017). We sequenced strand-specific RNA-seq libraries from ovaries of four homozygous wild-type adult females at 8mpf and testes from three homozygous wild-type adult male siblings, all of which were siblings of *amh^−26^* mutants. DESeq2 analysis showed that 16,493 genes were differentially expressed in wild-type adult ovaries vs. testes (Supplemental Table S2).

Principal component analysis separated adult testes and ovaries into two distinct groups widely separated in the PC1 axis, which explained 97% of the variance (Fig. 7A). Wild-type gonads separated from *amh* mutant gonads in the PC2 axis, which explained only 1% of the variance. Importantly, *amh* mutant ovaries tended to occupy the negative portion of the space and wild-type ovaries the positive portion, but the reverse was true for testes (Fig. 7A). This result shows that along the PC2 axis, the transcriptomes of mutant ovaries tended to be more like those of wild-type males (i.e., ovaries were masculinized) but the transcriptomes of mutant testes were more like female transcriptomes (i.e., testes were feminized). Masculinization of the ovary transcriptome and feminization of the testis transcriptome reflects the dual roles of *amh* in males and females.

**Figure 7.**
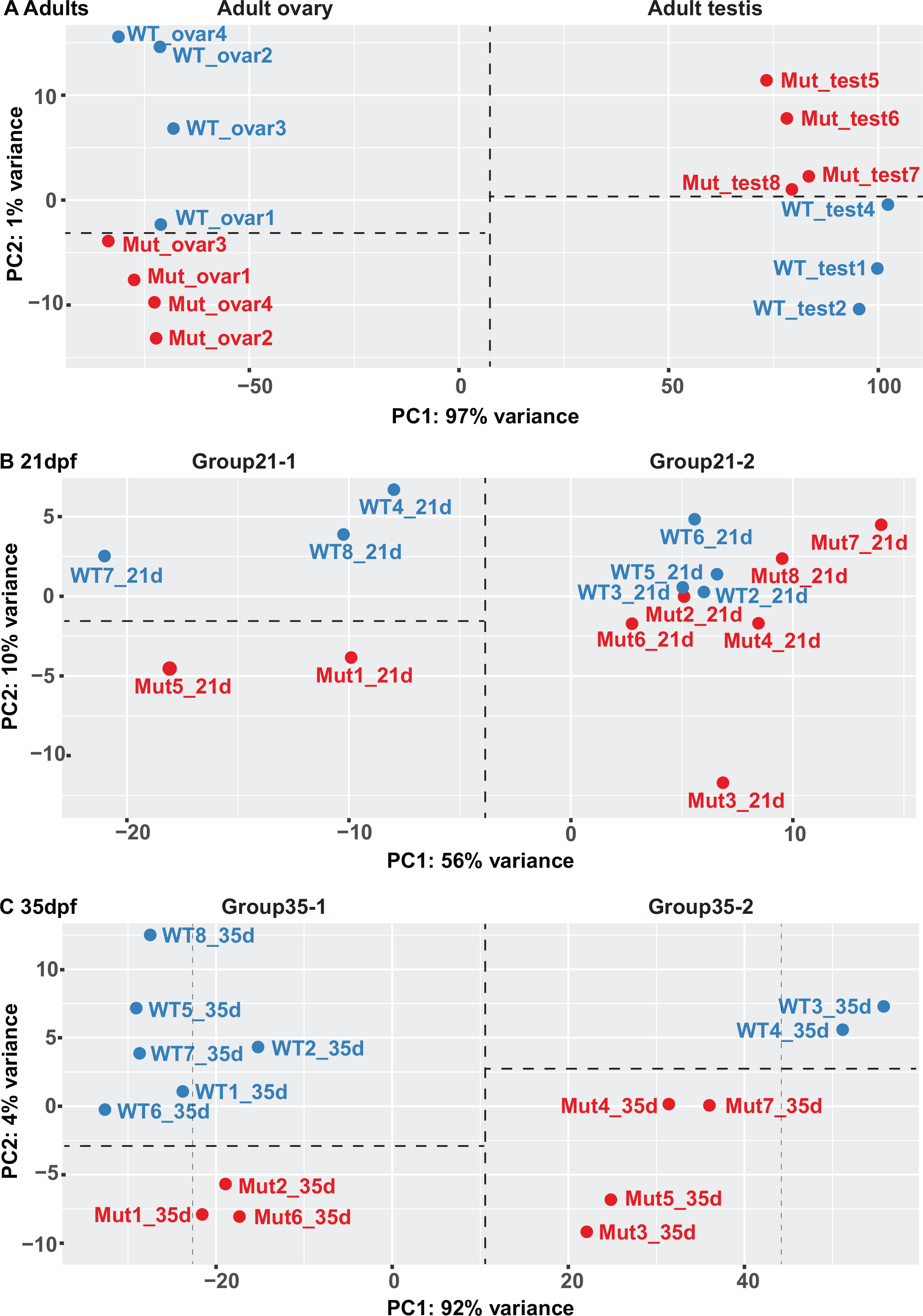
Principal Component Analyses (PCA). DESeq2-generated rlogs of the 500 most variable genes of (A) adult ovary and testes samples, (B) 21dpf samples, and (C) 35dpf samples.

Genes with the highest over-expression in adult zebrafish wild-type ovaries vs. wild-type testes tended to have no human orthologs and no previously assigned functions. For example, three genes were massively up-regulated in zebrafish ovaries with respect to testes (*zgc:171781, CABZ01059627.2, si:ch211-125e6.12*) by 146 million-, 120 million-, and 116-million-fold, respectively. Each of these three genes has several paralogs in zebrafish, but either no orthologs or few orthologs in other species and none have known functions, although ZFIN lists *si:ch211-125e6.12* as Pfam:PF00059, a C-type lectin. Of the 100 most up-regulated ovary genes, only 21 have gene names that imply function, including ten zona pellucida genes (*zp2.1, zp2.3, zp3.2, zpcx, zp2.5, zp2.6, zp2.2, zp3a.1, zp2l1, zp3a.2*), and only 11 other genes, including the ovary-specific epithelial cell tight junction gene *cldnd* (1582-fold up) (Clelland and Kelly 2011), the ovary-specific retinol saturase gene *retsatl* (1393-fold up) (Sreenivasan *et al*. 2008), the ovary carbonic anhydrase gene *ca15b* (1337-fold up) (Wang *et al*. 2013), the primordial germ cell histone gene *h1m* (1213-fold up) (Muller *et al*. 2002), two copies of the quinoid dihydropteridine reductase gene *qdprb2* (1079- and 490-fold up), the zebrafish ortholog of a gonadal soma nuclear repressor gene required for germ cell development *zglp1* (785-fold up) (Li S1 *et al*. 2007), the oocyte gene *cth1* (*cysteine three histidine 1*, 579-fold up (Tekronnie *et al*. 1999)), the germ plasm aggregation gene *birc5b* (510-fold up) (Nair *et al*. 2013), the extracellular matrix protein gene *ecm1a* (454-fold up), and the immune gene *crp2* (*C-reactive protein 2*, 409-fold up). We hypothesize that the large number of unannotated but highly expressed ovary-specific genes provide essential functions related to egg shells or other species-specific egg functions.

In addition to many genes of unknown function, most known female regulatory genes were also up-regulated in wild-type adult zebrafish ovaries compared to testes, including the Wnt-signaling genes *axin2* (24-fold up) and *rspo1* (2.1-fold up); the Foxl2-related genes *foxl2a* (ENSDARG00000042180, 49-fold up); *foxl2b* (ENSDARG00000068417, 7.4-fold up); and *foxl3* (ENSDARG00000008010, 5.3-fold up); the zona pellucida gene regulator *figla* (14-fold up), and other oocyte gene regulators like *bmp15* (41-fold up) and *gdf9* (26-fold up).

### Genome-wide transcriptomics of wild-type adult zebrafish testes

Up-regulated genes in wild-type testes vs. wild-type ovaries included the sperm-specific potassium ion channel gene *cngk* (9643-fold up) (Fechner *et al*. 2015). Genes encoding likely sperm components were the next most strongly over-expressed genes in adult wild-type testes vs. ovaries, including *ribc1* and *ribc2* (6094- and 3730-fold up, respectively), *ccdc83* (5898-fold up), and *rsph4a* and *rsph9* (5211- and 3569-fold up). Many genes annotated as being male-specific regulatory genes were also over-expressed in wild-type testis vs. wild-type ovary, including *amh* (244-fold up), *dmrt1* (411-fold up), *gsdf* (40-fold up), SoxD-related genes (*sox9a,* 47-fold up; *sox8a,* 14-fold up; *sox8b,* 27-fold up; *sox10,* 4.1-fold up), and *dhh* and its receptor-encoding genes *ptch1* and *ptch2* (61-, 3.9-, and 2.9-fold up in testes, respectively). The *wt1a* and *nr0b1* (*dax1*) genes were only slightly, but significantly, elevated in wild-type testes vs. ovaries (1.7-fold and 3.1-fold, respectively). Although vitellogenin genes appeared to be upregulated in wild-type testes vs. wild-type ovaries, overall counts were so low that fold changes were likely spurious. Vtg peptides have been detected in ovaries (Groh *et al*. 2013), although we saw no reads from *vtg* genes in wild type adult ovaries.

### Expression of steroid biosynthetic genes in wild-type gonads

Several steroid biosynthetic genes were differentially expressed comparing adult wild-type ovaries to wild-type testes. A duplication event in the zebrafish lineage after it diverged from *Astyanax* cavefish produced tandem co-orthologs of the single-copy human gene *CYP11A1*, which encodes side-chain cleavage enzyme, the first enzyme in steroid biogenesis. The *cyp11a1* gene was 25-fold up-regulated in zebrafish ovaries but *cyp11a2* was 8.1-fold up-regulated in testes, suggesting a subfunctionalization event (Force *et al*. 1999). The gene encoding Hsd17b1, which converts androstenedione to testosterone and estrone (E1) to estradiol (E2), was up-regulated in ovaries 175-fold over testes. Females convert testosterone to estrogen by aromatase, and *cyp19a1a* was up regulated 65-fold in wild-type ovaries compared to wild-type testes. Male mammals and male fish convert testosterone to 11-keto-testosterone, the primary androgen in fish, using Cyp11b1 in mouse (Cyp11c1 in zebrafish) and Hsd11b2 (Wang and Orban 2007; Yazawa *et al*. 2008; Lee *et al*. 2017); *cyp11c1* was up-regulated 1504-fold and *hsd11b2* was up-regulated 10.1-fold in wild-type testes vs. ovaries. *HSD3B1* and *HSD3B2* reside in tandem in human but their zebrafish orthologs are on two different chromosomes; we found *hsd3b2* up-regulated in ovaries (8.1-fold) and *hsd3b1* up-regulated in testes (6.4-fold).

This dataset (Table S2) contributes a substantial resource for understanding the normal functioning of adult zebrafish gonads and a standard for detecting the effects of mutations on gonad development.

### Gene expression in 21dpf zebrafish juveniles

At 21dpf, zebrafish late larvae are transitioning to become males or females (Takahashi 1977; Maack 2003; Rodriguez-MARI *et al*. 2005; Wang *et al*. 2007). Sequencing the gonad-containing trunks of *amh* mutants and wild types produced 189 million paired-end reads and after preprocessing (see Supp. Data Table 1), 103 million reads mapped to protein-coding exons. Analysis identified just 24 genes differentially expressed between *amh* mutants and wild types at 21dpf. The *amh* gene itself was under-expressed 6.1-fold in *amh* mutants, but this change was just outside the limit of significance (padj= 0.106), a result that reflects the relatively small difference between mutant and wild-type gonadal phenotypes as revealed by histology at this stage (see Fig. 3A-D) and the relative stability of transcripts from the mutated *amh* allele.

Transitional stage *amh* mutant fish at 21dpf expressed a number of gonadal regulatory genes abnormally. The most up-regulated gene in 21dpf *amh* mutant trunks vs. wild-type trunks was *nr0b2a* (3.03-fold up-regulated). In mammals, Nr0b2(SHP) dimerizes with Nr0b1(DAX1), thereby repressing Nr5a1(SF-1)-mediated activity of the *Amh* promoter (Tremblay and Viger 2001; Iyer *et al*. 2006). Furthermore, the loss of *nr0b1*(*dax1*) in zebrafish causes female-to-male sex reversal (Chen *et al*. 2016), in agreement with the reverse situation in which the duplication of *NR0B1* in humans causes male-to-female sex reversal (Barbaro *et al*. 2007). The up-regulation of *nr0b2a* in trunks of 21dpf *amh* mutants, as well as in adult mutant ovaries vs. wild-type ovaries (4.8-fold) suggests that *amh* normally represses *nr0b2,* and hence female development, in zebrafish. The second most up-regulated gene in 21dpf mutant trunks vs. 21dpf wild-type trunks, was the Leydig cell marker gene *cyp26a1* (2.95-fold up)(Wang *et al*. 2007), which encodes an enzyme that in zebrafish degrades retinoic acid (Rodriguez-MARI *et al*. 2013), the signal for entry into meiosis (Koubova *et al*. 2006; Adolfi *et al*. 2016). The up-regulation of *cyp26a1* in *amh* mutants would likely decrease the level of retinoic acid in mutants, and thus decrease the number of cells entering meiosis, a process that oocytes begin before spermatocytes do, thus suggesting that *amh* normally depresses *cyp26a1* expression at 21dpf. Other up-regulated genes in 21dpf *amh* mutants included the proteasome activator *psme4a* (2.4-fold up in mutants and 10.3-fold up in wild-type testes vs. wild-type ovaries); the lipid metabolism gene *trim63a* (2.1-fold up); the circadian nuclear receptor gene *nr1d1* (1.9-fold up in 21dpf *amh* mutants and 6.4-fold up in wild-type testes vs. wild-type ovaries); and the theca cell/Leydig cell marker *ptch2* (1.6-fold up in 21dpf *amh* mutants and 2.9-fold up in wild-type testes vs. wild-type ovaries) (Yao *et al*. 2002; Wijgerde *et al*. 2005; Herpin *et al*. 2013). Reciprocally, the most down-regulated gene in 21dpf mutant trunks vs. *amh* wild-type trunks was the complement factor H-related gene *cfhl1* (25.1-fold down in 21dpf mutant trunks and 25-fold down in wild-type ovaries vs. wild-type testes). Only two other genes were significantly down-regulated by more than two-fold in mutants: an uncharacterized sulfotransferase gene (*si:dkey-236e20.3*), and a hydroxybutyrate transporter gene *slc16a6b* (Hugo *et al*. 2012). We conclude that during the transitional period, the loss of *amh* function disrupts gonad development but not in a way that appears to strictly down-regulate canonical male-related genes as expected by the hypothesis that Amh should up-regulate male development.

Unsupervised similarity clustering split the fifteen 21dpf animals into two groups (Fig. 7B): Group21-1 contained two mutants and three wild types and Group21-2 had six mutants and four wild types. Principal component analysis (Fig. 7B) clustered individuals as they had with rlog-transformed Euclidean distances (Fig. 6), bolstering the view that these are biologically meaningful groups. The two groups are separated in the PC1 dimension, which explains 56% of the variance. PC2, which explains 10% of the variance, appeared to further separate Group21-1 into two groups: *amh* mutants and wild types, but small sample size thwarted statistical analysis of genes differentially expressed between Group21-1 *amh* mutants and wild types. The finding that Group21-1 and Group21-2 both contain wild-type and mutant individuals shows that at this early stage, *amh* expression is not the main factor that allocates individual fish into two groups.

To identify biological factors that distinguish the two synthetic 21dpf groups, we searched for genes differentially expressed between them. Analysis identified 440 genes that met the padj <0.1 criterion for false discovery rate (Supplemental Data Table S1). Genes up-regulated in Group21-1 vs. Group21-2 included several genes encoding components of the chorion, which oocytes begin to produce in stage IB follicles (Selman *et al*. 1993). These genes included the zona pellucida genes *zp2.2* (116-fold up-regulated in Group21-1 vs. Group21-2 and up-regulated 938-fold in wild-type ovary vs. wild-type testis), *zp2.5* (98-fold up and 958-fold up in wild-type ovary vs. wild-type testis), and 13 other *zp* genes. Zona pellucida genes in mouse and likely in zebrafish are controlled by the germ-cell transcription factor gene *figla* (*factor in germline-alpha*) (Liang *et al*. 1997; Onichtchouk *et al*. 2003; Mold *et al*. 2009); consistent with this role, *figla* was up-regulated in Group21-1 vs. Group21-2 (32-fold; 14-fold up in wild-type ovary vs. wild-type testis), Group21-1 also up-regulated the follicle stage I and II tight junction gene *cldnd* (96-fold; 5.6-fold up in wild-type ovary vs. wild-type testis), and other oocyte genes like the oocyte carbonic anhydrase gene *ca15b* (72-fold up; 1337-fold up in wild-type ovary vs. wild-type testis) (Wang *et al*. 2013), *zar1* (66-fold up; 148-fold up in wild-type ovary vs. wild-type testis) (Miao *et al*. 2017), *gdf9* (18-fold up; 26-fold up in wild-type ovary vs. wild-type testis), and *dazl* (11-fold up; not differentially expressed in wild-type ovary vs. wild-type testis) (Howley and Ho 2000; Clelland and Kelly 2011; Dranow *et al*. 2016). Germ cells in Group21-1 gonads were apparently entering meiosis because they up-regulated the synaptonemal complex gene *sycp2l* relative to Group21-2 (14.6-fold up; 14-fold up in wild-type ovary vs. wild-type testis). The strong expression of many oocyte genes shows that Group21-1 juveniles had substantially more developing oocytes than Group21-2. Vitellogenin genes were also up-regulated in Group21-1 trunks relative to Group21-2 trunks, including *vtg4* (32-fold up) and *vtg2* (31-fold). Vitellogenin genes were most likely expressed in liver, which was present in trunk preparations, but might also have been expressed in adipose cells in the ovary (Wang *et al*. 2005). This result suggests that Group21-1 gonads were already secreting estrogen that up-regulated *vtg* expression, but the only granulosa or theca cell marker that was up-regulated in Group21-1 compared to Group21-2 was *cyp11a1* (27-fold up), which encodes the enzyme catalyzing the first and rate-limiting step in steroid biogenesis. We conclude that genes over-expressed in Group21-1 vs. Group21-2 characterize developing oocytes.

Reciprocally, Group21-2 up-regulated 18 genes relative to Group21-1 fish (padj<0.1). Of these 18 genes, 14 were also up-regulated in wild-type testes relative to wild-type ovaries (*cap2, stard13a, si:ch211-133n4.4, col15a1b, adamts12, mmp13b, elf3, ift74, mhc1uka, b3gat1b, cyp27b1, BX004785.2, si:ch211-286b4.4, gstm.2*) an average of 49-fold; none were down-regulated in wild-type testis relative to wild-type ovaries; and four were not differentially expressed in wild-type gonads (*pomk, si:ch211-226h7.5, BX005421.3, zgc:162154*). We conclude that Group21-2, which was not expressing female genes, were expressing male genes, although few of these genes had previously been recognized as testis-related genes. Note, however, that the three most up-regulated genes in Group21-2 relative to Group 21-1 (*si:ch211-226h7.5, BX005421.3, zgc:162154*) were not differentially expressed in our ovary-vs.-testis comparison, suggesting that they may be transiently expressed in zebrafish transitioning to stable male development. We conclude that Group21-2 fish were embarking on a male pathway or were developmentally delayed with respect to Group21-1 fish.

We assessed the functional significance of the 440 genes differentially expressed between Group21-1 and Group21-2 samples using gene ontology (GO) analysis of biological processes (Mi *et al*. 2013). GO analysis identified 18 gene clusters at a false discovery rate (FDR) < 0.05. The top three clusters were strongly influenced by germ cell development. The highest loading enrichment cluster was “piRNA metabolic process” (FDR = 8.70E-03), and contained three genes (*henmt1, pld6,* and *asz1*) that were up-regulated in Group21-1 (putative females). The next highest loading enrichment cluster was “positive regulation of acrosome reaction,” (FDR 1.7E-12), with eleven of twelve genes annotated as zona pellucida genes or containing a zona pellucida domain. All were upregulated in Group21-1 samples. These same 12 genes were also the basis for the third (“egg coat formation”) and fourth (“binding of sperm to zona pellucida”) enrichment clusters. Together, the examination of individual dysregulated genes and the unbiased GO analysis agree that that among the 21dpf fish, Group21-1 juveniles are embarking on a female development and Group21-2 are becoming males. This is the first demonstration of a difference between developing males and females at this early age by whole-genome transcriptomic analysis.

### Gonadal gene expression in 35dpf juveniles

Sequencing the individual trunks of 15 juveniles at 35dpf (seven *amh^−28^* mutants and eight wild-type siblings) produced 93 million paired-end reads and after preprocessing (see Supplemental Data Table 2), 48 million reads mapped to protein-coding exons. Analysis of differential expression between wild-type and *amh* mutant samples identified 75 differentially expressed genes (Supplemental Data Table S1). Unlike the 21dpf late larvae, 35dpf mutant juveniles showed significant down-regulation of *amh* expression, the fourth most down-regulated gene in mutants (14.1-fold down).

The most differentially expressed up-regulated gene in 35dpf *amh* mutants was the butyrophilin subfamily immunoregulator gene *si:dkey-208m12.2* (85-fold up in mutants). Other strongly up-regulated genes in *amh* mutants were also immune related, including the novel fish interferon-stimulated genes *gig2l* (22-fold up) (Zhang *et al*. 2013), interferon-stimulated gene-15 (*isg15,* 4.8-fold up), the interferon-induced genes *mxe* (4.1-fold up) and *mxb* (5.0-fold up) (Novel *et al*. 2013). Interferon regulatory factor-7 (*irf7,* 4.5-fold up) is positively correlated with male-related genes in turbot (Ribas *et al*. 2016) and is a paralog of the trout master sex-determining gene *sdY*, a duplicated, truncated copy of *irf9* (Yano *et al*. 2012). Down-regulated genes in 35dpf *amh* mutants vs. wild-type siblings included the complement factor genes *cfhl2* (18-fold down, in *amh* 35dpf mutant trunks and 5.1-fold up in wild-type testes vs. wild-type ovaries) and *cfhl1* (18.7-fold down in *amh* 35dpf mutant trunks and 25-fold up in wild-type mature testes vs. ovaries); *cfhl1* was also the most strongly down-regulated gene in 21dpf *amh* mutant trunks vs. wild-type trunks (25.1-fold down). These results suggest that at 35dpf, gonads developing in *amh* mutants may experience cell damage that evokes an inflammatory response.

Similarity clustering based on globally correlated gene expression patterns resolved 35dpf samples into two distinct groups, and within those two major groups, wild types separated from *amh* mutants but with short branches in the tree (Fig. 6). Principal component analysis sorted 35dpf animals into the same two groups (Group35-1 and Group35-2), primarily along PC1, which explained 92% of the variance (Fig. 7C). Because each synthetic group included both *amh* mutants and wild types, differences other than genotype at the *amh* locus were important for distinguishing between major groups at 35dp. Within each of the two groups separated along PC1, mutants tended to occupy the lower portion of the plot and wild types the upper portion along the PC2 axis (Fig. 7C), even though this axis explained only 4% of the overall variance. Separation along PC2 may have resulted from expression changes in genes downstream of Amh function.

Analysis of genes differentially expressed between these Group35-1 and Group35-2 yielded 8,728 differentially expressed genes (Supplemental Table S1). The most differentially expressed genes between the trunks of Group35-1 and Group35-2 juveniles encode the egg yolk protein Vitellogenin-1 (*vtg1,* 929-fold up-regulated in Group35-1), with other *vtg* genes also highly upregulated (e.g., *vtg2,* 445-fold up in Group35-1 and *vtg4*, 319-fold up). The strong up-regulation of *vtg* gene expression in Group35-1 animals suggests first, that they are developing as females and second, that their livers had activated *vtg* genes due to secretion of higher levels of estrogen than Group35-2 fish, and thus, third, that their granulosa and theca cells were already functioning. Group35-1 animals also expressed differentially the female-enriched cell-cycle gene *btg4* (Small *et al*. 2009) (431-fold up), as well as several zona pellucida-encoding genes including *zp2l1* (247-fold up), *zpcx* (265-fold up), and *zp2.2* (214-fold up) along with their putative regulator *figla* (165-fold up). Group35-1 animals also expressed the meiosis gene *sycp2l* (120-fold up). These results show that Group35-1 animals had initiated a female pattern of developmental gene expression.

Reciprocally, the most up-regulated gene in Group35-2 relative to Group35-1 was transglutaminase-1-like-2 (*tgm1l2*, 65-fold up), which has not previously been documented as sex specific and has an unclear human ortholog, but was greatly over-expressed in wild-type testes vs. wild-type ovaries (72-fold up). This finding suggests the hypothesis that Group35-2 fish were embarking on a male developmental pathway. Group35-2 had increased expression of a number of other male-specific genes relative to Group35-1, including *amh* (46-fold up); the sperm-specific potassium ion channel gene *cngk* (18-fold up) (Fechner *et al*. 2015); an acyl-CoA thioesterase gene *acot17* (17-fold up) that was also over-expressed by adult wild-type testes vs. wild type ovaries (6.3-fold over-expressed in testes), but whose expression is otherwise unstudied; *ankar*, which human testes over-express compared to any other organ (Fagerberg *et al*. 2014) (14-fold up in Group35-2); the male factor *dmrt1* (13-fold up) (Webster *et al*. 2017); heat shock transcription factor 5 (*hsf5,* 11-fold up), whose human ortholog is expressed almost exclusively in testis (Fagerberg *et al*. 2014); *fank1*, the mammalian ortholog of which is exclusively expressed in pachytene spermatocytes and spermatids (Zheng *et al*. 2007) (11-fold up); the sperm-motility gene *t-complex-associated-testis-expressed-1* (*tcte1*, 9.8-fold up); and the testosterone-synthesizing enzyme gene *cyp11c1* (8.2-fold up in Group35-2). These results show that Group35-2 individuals were becoming males. Group35-2 individuals also had increased expression of the synaptonemal complex encoding genes *sycp3* (16-fold up in Group35-2) and *sycp2* (8.7-fold up), and the DNA meiotic recombinase-1 gene (*dmc1*, 11-fold up), likely reflecting a large number of spermatogonia preparing to undergo meiosis in Group35-2 fish compared to fewer meiotic cells in Group35-1 individuals. We conclude that Group35-2 fish were beginning to mature their testes, as judged by their stronger expression of male-related genes compared to Group35-1 (putative females).

Within Group35-2, all of the *amh* mutant samples were substantially shifted in the PC1 dimension towards Group35-1 with respect to the wild-type samples. Because Group35-1 were expressing female genes and Group35-2 were expressing male genes, this finding shows that 35dpf fish lacking *amh* activity tend to be feminized in terms of their gene expression. Likewise, within Group35-1, all three of the *amh* mutants were closer to Group35-2 than five of the six wild types in the PC1 axis. This result suggests that zebrafish juveniles developing as females tend to be somewhat masculinized in the absence of *amh* activity. These observations confirm the utility of Amh in both male and female development.

Expression patterns of the 35dpf putatively male (Group35-2) fish were strongly correlated to expression patterns of the 21dpf Group21-2 (not obviously female) fish. Of the 18 genes that were significantly differentially up-regulated in putative non-female Group21-2 fish (Fig. 7C), eleven (*zgc:162154, BX005421.3, BX004785.2, mhc1uka, elf3, adamts12, mmp13b, cap2, stard13a, si:ch211-133n4.4, col15a1b*) were also significantly differentially up-regulated in the male gene-expressing 35dpf cohort (Group35-2, Fig. 7C) at an average of 2.9-fold, with the amount of up-regulation highly correlated between the 21dpf and 35dpf datasets (correlation coefficient of 0.97). The seven other genes significantly up-regulated in the not-female Group21-2 (*pomk, ift74, b3gat1b, cyp27b1, si:ch211-286b4.4, gstm.2, si:ch211-226h7.5*) were not differentially expressed between the two 35dpf synthetic groups. Of the 25 most up-regulated differentially expressed genes in the male-like Group35-2 relative to Group35-1, all but one were also up-regulated in wild-type testis relative to wild-type ovary an average of 1922-fold (*tgm1l2, zgc:158427, amh, cngk, acot17, sycp3, gstk4, ankar, dmrt1, si:ch211-242f23.3, pimr214, hormad1, si:dkeyp-50b9.1, ifit16, hsf5, dmc1, fank1, si:dkeyp-80c12.8, ttc29, spag16, tcte1, dnah6, hbaa2, tekt1*). Only one gene up-regulated in Group35-2 (*CABZ01076758.1*) was not differentially expressed in wild-type testis compared to wild-type ovary. We conclude that, despite the fact that up-regulated genes in the non-female 21dpf group were mostly not previously known to be male-related genes, their continued up-regulation in the group of 35dpf fish that were expressing many clearly male genes shows that fish in Group21-2 were also developing male characteristics. These experiments thus identify a previously unknown cohort of sex-specific genes expressed early in gonadogenesis.

Gene ontology analysis of differentially expressed genes comparing the two 35dpf groups yielded 44 enrichment clusters (FDR, p<0.05). The most significantly enriched cluster contained 22 genes enriched for “negative regulation of mitotic cell cycle phase transition” (FDR=3.0E-02). Among these genes were mitotic checkpoint genes (*bub1bb, bub1, hus1, mad1l1, and rad17*), and a variety of DNA repair genes (*oraov1, orc1, mre11a, blm, and msh6*). All were up-regulated in female-like Group35-1. The second cluster included 30 genes enriched for “mitotic cell cycle checkpoint” (FDR= 5.93E-03), with an expanded list of checkpoint and DNA repair genes similar to the first cluster, including *rad9a, rad9b, eme1,* and *msh2*. All were up-regulated in Group35-1 except *rad9b*, suggesting a negative correlation of the co-orthologs *rad9a* and *rad9b* and their possible subfunctionalization. The third enrichment cluster comprised 41 genes enriched for “DNA-dependent DNA replication” (FDR = 3.60E-03). These included a variety of DNA polymerases (*polg, poln, pola2, pole2,* and *pold2*) and associated DNA binding proteins (*orc1, orc3, orc6, rpa1, msh2, cdc45,* and *wdhd1*). All but *poln* were up-regulated in Group35-1, female-like fish. This cluster also included the up-regulated early onset breast cancer and Fanconi anemia gene *brca2*(*fancd1*). Nine other Fanconi anemia genes were also significantly up-regulated in the DESeq2 analysis for Group35-1 vs. Group35-2 (*fanca, fancb, fancc, fancd2, fance, fancf, fancg, fanci, fancm*). These GO enrichment terms for the two 35dpf juvenile groups differed markedly from the GO terms discovered for the two 21dpf late larval groups. At 21dpf, germ cell functions (piRNAs and egg-shell genes) dominated GO terms differentially expressed between the two groups, but at 35dpf, cell cycle and DNA-repair genes were most differentially expressed between the two groups.

### Amh activities regulate adult testis gene expression patterns

To help understand the molecular genetic basis for abnormal testis morphologies caused by loss of *amh* function, we sequenced seven libraries of 8mpf adult testes, one each for three wild-types and four *amh^−26^* mutants. Sequencing produced 54 million paired end reads, of which 24 million passed quality filters and mapped to the zebrafish genome assembly. DESeq2 identified 3,902 differentially expressed genes (Supplemental Table S2). Similarity clustering (Fig. 6) and PCA based on correlated gene expression (Fig. 7A) placed all seven testis samples together, with clear separation between wild types and *amh* mutants. As predicted, *amh* was significantly down-regulated in mutant testes (*amh* 6.7-fold down in mutants).

The first three most up-regulated genes in mutant testes vs. wild-type testes were the same as the first three most up-regulated genes in wild-type ovaries vs. wild-type testes (*CABZ01059627.2, si:ch211-125e6.12, zgc:171781,* up-regulated about 239,000-, 116,000-, and 90,000-fold respectively). This result shows that *amh* mutant testes greatly up-regulated ovary-specific genes, suggesting a partial feminization of adult *amh* mutant testes, which could happen by the retention of the early oocytes that were detected by histology (see Fig. 4O, S).

Leydig cell markers were mostly down-regulated in *amh* mutant testes compared to wild-type testes, consistent with our previous analysis of *nr5a1a* expression, which labels Leydig cells (Fig. S, S’). Zebrafish orthologs of 16 of 50 Leydig cell marker genes (Uhlen *et al*. 2015) were differentially expressed in zebrafish *amh* mutant testes vs. wild-type testes. The zebrafish orthologs of 14 of these 16 human Leydig cell marker genes (*DHH, APOE, AMH, FDX1, CYP17A1, AK1, CNTRL, SGPL1, EPHX1, CCS, ELOA2, ACLY, TMF1, CLEC16A*) were down-regulated in *amh*-mutant testes an average of 3.6-fold, and orthologs of only two (*CACNA1H* and *PRSS12*) were up-regulated (average of 2.8-fold). The down-regulated genes included *dhh* (down 13-fold), which triggers Leydig cell differentiation (Yao *et al*. 2002), and *cyp17a1* (down 4.3-fold), which encodes the second enzyme in the testosterone biosynthesis pathway. We conclude that loss of *amh* function disrupts the specification, proliferation, or functioning of Leydig cells in adult zebrafish testis, supporting conclusions from our *in situ* hybridization analyses (see Fig. 5N, N′).

Sertoli cell marker genes responded in various ways to the loss of *amh* activity. Zebrafish has orthologs of 38 of 50 human Sertoli cell marker genes (Uhlen *et al*. 2015). Several Sertoli cell genes were up-regulated in mutant testis relative to wild-type testis (*gsdf*, 3.4-fold up; *fndc7a,* 2.40-fold; and *ddx3a*, 1.30-fold) but seven were down-regulated in mutants, including *amh* itself (2.8-fold down, as well as *arid4a, alg11, fndc3a, lrig1, abhd2b, sox8b,* and *cst3*, an average of 2.1-fold down (Supplemental Table 2). Many Sertoli cell regulatory genes, including *acvr2aa, acvr2ab, dmrt1, fshb, fshr, hsd17b4, inha, nr0b1, sox9a, and sox9b*, were not differentially expressed between mutant and wild type testes. These results suggest that *amh* tends to inhibit some Sertoli cell functions, but not strongly. Recall that our *in situ* hybridization data showed mainly a change in the spatial distribution of Sertoli cells (see Fig. 5B, B′, H, H′).

Germ cells in *amh* mutant testes seemed to develop rather normally from a histological perspective, although mutant testes accumulated many more germ cells than normal (see Fig. 4U, U’). The Human Protein Atlas lists 50 genes strongly expressed in spermatogonia, but only 15 of these have orthologs or closely related paralogs in zebrafish. None of these 15 germ cell genes was up-regulated in *amh* mutant testes vs. wild-type testes and only three (*brd2a, CU929144.1 (*alias *cfap46),* and *meiob*) were down-regulated (an average of 2.3-fold), while twelve (*acvr2ba, acvr2b, brip1, cbl, dazl, dmrt1, dync1h1, hmga2, mgea5, mgea5l, mtmr3, nanos3, plk4, uchl1*) were not differentially expressed. For 51 human spermatocyte marker genes in the Human Protein Atlas, only 19 have zebrafish orthologs. One gene (*BCL6*) has two zebrafish co-orthologs, and although one co-ortholog (*bcl6a*) was up-regulated (2.5-fold) in *amh* mutant testes, it’s co-ortholog (*bcl6b*) was down-regulated (5.5-fold). The zebrafish orthologs of seven human spermatocyte genes were down-regulated an average of 3.1-fold (*bcl6b, clgn, cremb, ccdc65, crema, rnf32, tekt1*) consistent with a role for *amh* in spermatogenesis.

Gene ontology analysis of *amh* mutant testes vs. wild-type testes identified ten GO clusters (FDR<0.05). The highest loading cluster included 22 genes enriched for “interciliary transport,” including intraflagellar transport proteins (e.g. *ift27, ift57, ift74,* and *ift81,* all down-regulated in mutants 2- to 3-fold) and tetratricopeptide repeat domains (*ttc21b* and *ttc26* (1.9 and 2.1 fold down-regulated)). The second cluster contained 38 genes enriched for “axoneme assembly”, including dynein-related genes down-regulated in mutants (e.g. *dnah3, dnah5, dnah12, dnai1.2*), and genes encoding coiled-coil domain proteins (*ccdc39, ccdc103, ccdc114,* and *ccdc151*). Down-regulation of genes involved in microtubule assembly is consistent with defects in the ability of *amh* mutants to produce mature sperm with fully developed tails. We conclude that sperm maturation was suppressed in *amh* mutant testes, which likely contributed to observed loss of fertility as animals aged.

### Amh activities regulate adult ovary gene expression patterns

To understand the genetic effects of Amh on zebrafish ovary development, we investigated gene expression patterns of ovaries from four *amh* mutants and four of their wild-type siblings in eight individual libraries. Sequencing produced 114 million paired-end reads, 60 million of which mapped to the Ensembl v91 protein coding exons in GRCz10. Similarity clustering using the entire 45-sample dataset placed all ovary samples together on a long branch, indicating a unique transcriptional profile (Fig. 6). Within the eight ovary samples, *amh* mutant ovaries occupied a different branch from wild-type ovaries (Fig. 6); this result contrasts to the 21dpf and 35dpf samples where *amh* mutants and wild types intermixed within each age group and within sub-groups (Fig. 6). We conclude that developmental processes that depend on Amh are stronger in adult ovaries than in 21dpf or 35dpf animals.

DESeq2 identified 7,426 genes differentially expressed in mutant vs. wild-type ovaries (Supplemental Table S2). Although *amh* was down-regulated in mutant ovaries compared to wild-type ovaries (2.4-fold down), this difference was just short of reaching statistical significance (padj=0.102). A comparison of adult *amh* mutant ovaries to adult wild-type ovaries showed that the top six most differentially expressed genes in terms of fold change encode vitellogenins. Zebrafish express vitellogenin genes not only in their livers in response to estrogen as do egg-laying tetrapods, but also in adipocytes in their ovaries (Wang *et al*. 2005). Over-expression of *vtg* genes in *amh* mutant ovaries is not due to contamination from liver in our samples by dissection errors because zebrafish liver marker genes, such as *fga, fgb, fabp10a, hmgcra,* and *hmgcrb*, as well as zebrafish orthologs of human liver marker genes including *apoa2, a1bg, ahsg, f2, cfhr2, hpx, f9* (Uhlen *et al*. 2015) were not differentially expressed between mutant and wild-type ovaries in our samples. This result is consistent with our morphological studies, which showed that the mutant ovary accumulates enormous quantities of follicles stalled in a pre-vitellogenic state (Fig. 4E, F). If a negative feedback mechanism were in place that senses yolky oocytes and inhibits the transcription of *vtg* genes in ovarian cell types (Wang *et al*. 2005), then the absence of yolky oocytes would result in continuous up-regulation of *vtg* gene expression.

Many ovarian regulatory genes were greatly under-expressed in adult mutant ovaries compared to wild-type ovaries. Granulosa cell marker genes (Hatzirodos *et al*. 2015) tended to be down-regulated in *amh* mutant ovaries vs. wild-type ovaries, including aromatase (*cyp19a1a,* 18.7-fold down in mutant ovaries), *nr5a2* (12.4-fold down), luteinizing hormone receptor (*lhcgr,* 12.6-fold down), *gata4* (25.8-fold down), the estrogen receptors *esr1* (4.0-fold down) and *esr2b* (6.9-fold down), *foxl2a* (ENSDARG00000042180, 5.7-fold down), *foxl2b* (ENSDARG00000068417, 4.4-fold down), and *slc35g1* (7.3-fold down). Theca cell marker genes (Hatzirodos *et al*. 2015) were also down-regulated: *insl3* (23.6-fold down), *nid1b* (10.3-fold down), *nr5a1a* (16.1-fold down), *nr5a1b* (2.9-fold down), *star* (3.8-fold down), *cyp11a2* (3.4-fold down), and *hsd3b1* (3.4-fold down). These results show that expression of marker genes for both granulosa cells and theca cells are down-regulated and confirm *in situ* hybridization results (Fig. 5) that showed significant disruption of follicle cell morphologies in *amh* mutants.

Given the great enlargement of *amh* mutant gonads (Fig. 4A, B, E, F), it was unexpected to find that many key oocyte marker genes were not expressed differentially between *amh* mutants and wild types, including *vasa*, *dnd1, piwi* paralogs, *bmp15, gdf9*, *nanos* paralogs, *sycp* synaptonemal complex genes, and zona pellucida genes (Liu *et al*. 2006). This finding was further surprising given that most of these genes were mis-regulated in the putative female vs. putative male synthetic groups for the 21dpf and 35dpf time points. Some markers of meiosis were up-regulated in mutant ovaries compared to wild-type ovaries (*spo11* (2.1-fold up), *rad51d* (3.1-fold up)), but others, like *dmc1* and synaptonemal complex genes, were not differentially expressed. These results may suggest that the large number of mutant oocytes accumulated by mutant ovaries had stalled at an early stage of meiosis.

Gene ontology analysis (PANTHER (Mi *et al*. 2013)) identified 49 enrichment clusters comparing genes differentially expressed between adult *amh* mutant ovaries and adult wild-type ovaries (FDR<0.05). The highest loading cluster was “ribosomal large subunit assembly” (FDR = 2.59E-02), and was comprised of 17 genes, including various ribosomal protein genes (*rpl3, rpl5a, rpl5b, rpl6, rpl11, rpl12,* and *rpl23a*) that were up-regulated in *amh* mutant ovaries compared to wild-type ovaries. The second cluster was “cytoplasmic translation” (FDR = 1.49E-03), and was comprised of 28 genes including additional upregulated ribosomal protein genes (e.g. *rpl7, rpl9, rpl22l1, rpl26, rpl29, and rpl31*), and translation initiation factors (*etf1b, eif3a, eif3m, eif4h, and eif4bb*). The third cluster was “maturation of large subunit-rRNA (FDR = 4.81E-02) and contained 18 up-regulated ribosomal biogenesis protein genes (e.g., *wdr12, nsa2, las1l,* and *rpf2*), as well as additional ribosomal proteins (e.g. *nhp2, rpl7, rpl10a, and rpl35*). These clusters reflect the massive accumulation of ribosomes that maturing eggs normally store. Other gene ontology enrichments indicated coordination of replication, transcription, and translation. These results are expected from the morphological studies that showed massive changes in oocyte accumulation and defects in follicle development (Fig. 4E, F, I, J).

### The lack of *amhr2* in the genomes of zebrafish and other cyprinids is associated with a chromosome rearrangement breakpoint

AMH in mammals binds to the receptor AMHR2. Humans lacking function of either *AMH* or *AMHR2* have persistent Müllerian ducts and *Amh;Amhr2* double mutant mice have the same phenotype as either single mutant, showing that the ligand and receptor act in the same pathway (Imbeaud *et al*. 1996; Mishina *et al*. 1996). Amhr2 makes a dimer with one of the Bmpr1 proteins, and the zebrafish mutant phenotype for *bmpr1bb* mimics the *amh* mutant phenotype reported here for the enlarged testes and accumulation of immature oocytes, but the *bmpr1bb* mutant males did not retain oocytes like the *amh* mutant testes did (Neumann *et al*. 2011). In addition, the *bmpr1bb* mutants did not appear to alter the sex ratio as did the *amh* mutants (Neumann *et al*. 2011). Percomorph fish genomes generally contain an ortholog of *Amhr2* (e.g., stickleback, (*Gasterosteus aculeatus*, ENSGACG00000006672). In contrast to percomorphs, zebrafish is an otophysan teleost, and at least two suborders of otophysans also possess an *amhr2* gene, the characiform suborder, including both red-bellied piranha (*Pygocentrus nattereri*, ENSPNAG00000001197) and cavefish (*Astyanax mexicanus,* ENSAMXG00000024722), and the siluriform suborder, including channel catfish (*Ictalurus punctatus*, ENSIPUG00000006414). In contrast, zebrafish in the cypriniform suborder of otophysans appears to lack *amhr2* (Ribas *et al*. 2016).

To understand whether this apparent loss of *amhr2* is unique to zebrafish (which could then either be a zebrafish-specific loss or a genome assembly error) or whether it might represent an event shared among cypriniforms, we studied conserved syntenies. Results showed that the *amhr2-*containing region of cavefish (Supplemental Figure S2A) corresponds to three widely scattered portions of the zebrafish genome (Fig. S2B, C), with a break in conserved syntenies occurring at the predicted location of *amhr2* and its nearest neighbor (ENSAMXG00000024723, *cell division cycle associated 7*), which is also present in most percomorphs but is missing from zebrafish. Duplicates of the zebrafish orthologs of genes flanking *amhr2* in cavefish that originated in the teleost genome duplication are on two different zebrafish chromosomes, Dre2 and Dre22 (Fig. S2). These data are consistent with the hypothesis that both *amhr2* and its neighbor ENSAMXG00000024723 disappeared in the zebrafish lineage after it diverged from other otophysans associated with a chromosome inversion breakpoint at the ancestral site of *amhr2*.

To determine whether the rearrangement breakpoint at the expected position of *amhr2* is zebrafish specific, we examined the genomes of two other cypriniform fish: common carp (*Cyprinus carpio*) and goldfish (*Carassius auratus*). BLASTP searches using cavefish Amhr2 against common carp and goldfish genomes did not identify an Amhr2 ortholog, but brought back Tgfbr2b as the most similar protein, suggesting that these cyprinids, like zebrafish, have no ortholog of *amhr2*. (Note that BLASTP searches of cavefish Amhr2 vs. zebrafish brought back as the two top hits Bmpr2a and then Bmpr2b.) Conserved synteny analysis showed that zebrafish and common carp share gene orders at the location predicted for *amhr2* (Supplemental Fig. S2). This result is predicted by the hypothesis that a chromosome rearrangement with a breakpoint in or near *amhr2* destroyed this gene and that the event occurred after cyprinid otophysans diverged from characiform and siluriform otophysan teleosts, but before the divergence of the zebrafish and carp lineages.

## DISCUSSION

In human male fetuses, Sertoli cells secrete Anti-Müllerian Hormone, which causes developing Müllerian ducts to disappear, while in adult human females, AMH suppresses the initiation of primary follicle growth, serves as a marker for ovarian reserve, and provides an assay for conditions like polycystic ovarian syndrome (PCOS) (Carlsson *et al*. 2006; Diamanti-KANDARAKIS 2008). Teleost fish do not have Müllerian ducts but nevertheless maintain an *amh* gene (Adolfi *et al*. 2018). To investigate conserved roles of Amh in vertebrates that lack a Müllerian duct, we made zebrafish *amh* null activity alleles.

Results showed that *amh* activity promotes, but is not essential for, male development in zebrafish because homozygous *amh* mutants were only about 20% as likely to develop into males as their wild-type siblings (see also (Lin *et al*. 2017)). In addition, most mature adult male mutant gonads we examined contained a few early stage oocytes while at the same age, wild-type siblings did not. This finding shows that Amh normally masculinizes zebrafish by inhibiting the development or survival of young oocytes. Several fish species expand the male-biasing role of *amh,* having evolved a modified gene duplicate that has become the major sex determinant. For example, Patagonian pejerrey and Nile tilapia possess *amh* gene duplicates that independently became the primary sex determinant (Hattori *et al*. 2012; Li *et al*. 2015); ling cod also possesses a male-specific *amh* duplication (Rondeau *et al*. 2016). A further demonstration of the role of the *amh* pathway is the finding that variants of the Amh receptor gene *amhr2* provide the primary sex determinant in several species of pufferfish (Kamiya *et al*. 2012; Ieda *et al*. 2018). In tetrapods and most teleosts, Amh receptor type II (Amhr2) is expressed in Leydig and Sertoli cells (Racine C *et al*. 199; Di Clemente N *et al*. 1994) and mediates Amh signaling. Zebrafish, however, has no ortholog of *Amhr2*, which we show here is associated with an inversion breakpoint. Because we show that Amh is critical for zebrafish gonad development, the function of Amhr2 is likely performed by another Bmpr2 paralog. Zebrafish has two *bmpr2* ohnologs; *bmpr2a* is expressed in young oocytes and in ovarian follicle cells and *bmpr2b* is expressed in follicle cells (Li and Ge 2011; Dranow *et al*. 2016). We found that *bmpr2a* was over-expressed in adult wild-type testis vs. wild-type ovary (4.7-fold), and both *bmpr2a* and *bmpr2b* were under-expressed in *amh* mutant ovary vs. wild-type ovary (4.7-fold and 6.2-fold, respectively), but we did not detect differential expression of any *bmpr2* genes comparing putative female and putative male groups of 21dpf or 35dpf juveniles.

Young zebrafish *amh* mutant females are fertile, showing that they have functional reproductive ducts. As female *amh* mutants age, however, they become sterile, showing that Amh supports continued female fertility. Zebrafish males lacking *amh* activity are less effective than their wild-type male siblings at stimulating wild-type females to lay eggs, showing that Amh action improves male mating behavior. It is as yet unknown whether this difference is related to changes in brain organization that depend on the developmental availability of Amh. Likewise, mouse mutants lacking either Amh or Amhr2 show feminized spinal motor neurons and some feminized behaviors (Wang *et al*. 2009). Some wild-type eggs that were fertilized by mutant males develop to hatching, showing that functional male reproductive ducts form without the benefit of Amh. Nevertheless, eggs laid by wild-type females in the presence of *amh* mutant males are less likely to develop than those fertilized by wild-type males, showing that Amh helps optimize sperm production, function, or release. We conclude that in zebrafish, Amh is not necessary for the development of reproductive ducts or for the initiation of functional gamete formation, but is necessary for continued fertility in both sexes.

Wild-type siblings of our *amh* mutants had gonads with stage I oocytes at 21dpf as expected (Takahashi 1977; Selman *et al*. 1993; Rodriguez-MARI *et al*. 2005; Wang and Orban 2007; Rodriguez-MARI *et al*. 2010), but some 21dpf *amh* mutants had gonads that contained only undifferentiated germ cells, suggesting that *amh* activity helps to accelerate gonad development. By 35dpf, about half of wild-type siblings were continuing to develop oocytes and the other half were forming spermatocytes, but most *amh* mutants were developing normal-looking ovaries and few were developing spermatogonia. We conclude that *amh* mutants are slow to adapt a male phenotype, and many never do, leading to a female-biased sex ratio among *amh* mutants.

Ovaries in mature adult *amh* mutant females were swollen by immature oocytes to nearly three times the size of ovaries in wild-type siblings, confirming previous results (Lin *et al*. 2017). The ovarian phenotype of zebrafish *amh* mutants mimics the phenotype of *gsdf* mutants in zebrafish (Yan *et al*. 2017), of Amh receptor mutants in female medaka (Hattori *et al*. 2012), and of polycystic ovarian syndrome in humans (Diamanti-KANDARAKIS 2008). We conclude that Amh represses primordial oocyte proliferation but stimulates oocyte maturation in zebrafish.

Suppression of oocyte maturation appears to be a role of Amh shared by fish and mammals because mice lacking Amh show premature depletion of the primordial follicle pool (Durlinger *et al*. 1999), likely because Amh slows follicle growth in zebrafish and mammals as it does in humans (Carlsson *et al*. 2006). Zebrafish males lacking *amh* activity also had greatly enlarged gonads (Lin *et al*. 2017), showing that in zebrafish, Amh helps slow gonad growth both in males and in females. In contrast to females, however, males even at 18mpf appeared to contain mature gametes, although these males were sterile, demonstrating -- somewhat paradoxically -- that, although Amh normally nudges juvenile zebrafish towards a male pathway, its activity in older fish is required for oocyte maturation (no *amh* mutant female produced mature eggs at 11mpf) but appears only to accelerate spermatocyte maturation (3% of eggs laid by wild-type females developed after mating with mutant males at 11mpf).

The finding of the similarity of the phenotypes of zebrafish *amh* and *gsdf* mutants raised the question of whether these genes regulate germ cell proliferation and differentiation by acting in the same or different pathways. Because results showed that *amh;gsdf* double mutant gonads were no more compromised than either single mutant, we conclude that *amh* and *gsdf* act in the same pathway. Sertoli cells and granulosa cells both express both *amh* and *gsdf* (Rodriguez-MARI *et al*. 2005; Von Hofsten *et al*. 2005; Gautier *et al*. 2011a; Gautier *et al*. 2011b; Yan *et al*. 2017). Zebrafish lacks an ortholog of *amhr2,* which encodes the Amh receptor found in other vertebrates, and the receptor for Gsdf is unknown, so the relative positions of Amh and Gsdf in a shared developmental pathway are currently ripe for further investigation.

Differences in gene expression patterns in mutants compared to wild types give clues to how genes exert their effects. We studied altered gene expression patterns in *amh* mutants in two ways: by *in situ* hybridization and by whole genome transcriptome analyses. *In situ* hybridization experiments revealed reduced *amh* transcript accumulation in granulosa cells and disrupted organization of *amh-*expressing Sertoli cells. The lack of Amh activity appeared to result in an up-regulation of *gsdf* expression in Sertoli cells both in our *in situ* hybridization data and in our RNA-seq data (3.39-fold up-regulated in mutant testes (Supplemental Table 2)), suggesting that if Amh and Gsdf act in the same pathway, Amh may repress *gsdf* activity. Reciprocally, *amh* expression was up-regulated 1.8 fold in our *gsdf* mutants (Yan *et al*. 2017), showing interdependent regulation of *amh* and *gsdf* genes. The down-regulation of the aromatase gene in zebrafish *amh* mutants and the failure of aromatase-expressing cells to completely envelop oocytes suggests that Amh is required for proper development of granulosa cells to the stage appropriate for maximal aromatase expression. With depressed aromatase activity, levels of estrogen should diminish, thereby inhibiting oocyte maturation, which we observed. An additional contributor to endocrine disruption would be our finding of the greatly reduced expression of *nr5a1a* – which encodes a nuclear receptor transcription factor that regulates steroidogenesis genes (Val *et al*. 2003). Our in situ hybridization results compared to histological phenotypes paint a picture of disrupted development stemming from *amh-*expressing granulosa and Sertoli in zebrafish *amh* mutants that leads to disordered organization of these helper cells, their failure to support sex steroid output of theca and Leydig cells, followed by failure of proper germ cell maturation and inhibition of germ cell proliferation.

Genome-wide transcriptional analyses further provided an unbiased probe of the mechanisms of normal gonad development and the roles of Amh. A comparison of wild-type ovary to wild-type testis identified hundreds of genes with previously unknown functions, many of which are lineage-specific, with strong differential expression. These genes likely encode egg shell and sperm components. These data provide, for the first time, information related to function for many genes known previously only by sequence.

Principal component analysis showed that the transcriptomes of adult *amh* mutant ovaries were shifted in the direction of wild-type testis transcriptomes and mutant adult testis transcriptomes were shifted in the direction of wild-type ovary transcriptomes. This result likely reflects the depressed ability of *amh* mutant ovaries to convert testosterone to estrogen and the observed retention of immature oocytes in adult *amh* mutant testes.

Analysis of the transcriptomes of gonad-containing trunks of juveniles provided important insights into both normal zebrafish sex determination and the roles of Amh in gonad development and physiology. At 21dpf, laboratory strain zebrafish have morphologically undifferentiated gonads (Takahashi 1977; Maack 2003; Rodriguez-Mari *et al*. 2005; Wang *et al*. 2007), so it was a surprise to find that unsupervised transcriptome similarity clustering divided late larvae into two distinct groups. Each of these two groups contained both *amh* mutants and wild types, showing that factors other than *amh* function distinguish these two groups. Differential expression analysis showed that one group over-expressed ovary genes and the other over-expressed genes that were also over-expressed in wild-type testes vs. wild-type ovaries, although few of these genes had previously been recognized as testis marker genes. We conclude that, despite little morphological differentiation in 21dpf gonads, they had already embarked on a female or male developmental program. This is the first report of the genome-wide transcriptional differentiation of zebrafish gonads at such an early age.

The small number of genes (16) that were differentially up-regulated in *amh* mutant trunks vs. wild-type trunks at 21dpf included regulators of steroidogenesis and meiosis. Two up-regulated genes in mutants affect development or function of steroid-producing Leydig cells: *nr0b2*, which inhibits steroidogenic gene expression in mouse Leydig cells (Volle *et al*. 2007), and *ptch2,* part of a receptor complex for desert hedgehog signaling in Leydig cells (Yao *et al*. 2002; Wijgerde *et al*. 2005; Herpin *et al*. 2013). Amh also likely helps regulate the initiation of meiosis. Nr0b2 in mouse reduces the level of retinoic acid, thereby reducing the expression of genes essential for mitotic germ cells to enter meiosis (Volle *et al*. 2007) and we found that *nr0b2* was the most up-regulated gene in 21dpf *amh* mutants. Late larval zebrafish *amh* mutants also up-regulated the Leydig cell marker gene *cyp26a1,* which encodes an enzyme that degrades retinoic acid, the regulator of entry into meiosis. We conclude that Amh from Sertoli cells is required for normal Leydig cell development and function and likely helps regulate the timing of meiosis as early as 21dpf.

Transcriptomes of 35dpf trunks also separated samples into two groups, each of which contained both mutants and wild types. Within each group, mutants separated from wild types, showing that between 21dpf and 35dpf, *amh* had begun to exert a significant effect on gonad development. One group evidently had significant levels of estrogen because they were strongly expressing estrogen-induced vitellogenin genes, and in addition they strongly expressed zona pellucida egg shell genes. The other group over-expressed a number of testis-specific genes including *amh* and *dmrt1* as well as genes not previously recognized as testis genes but over-expressed in wild type testis vs. wild-type ovary, thus providing novel insight into potential functions of these genes previously known only by sequence. Expression levels of genes in putatively male 35dpf animals were strongly correlated to their levels in the not-female 21dpf group, confirming that some late larval fish had already begun to initiate a male developmental pathway. GO enrichment terms between putative males and putative females in 35dpf juveniles changed from a focus on germ cell functions like piRNAs and egg-shell genes at 21dpd to an emphasis on cell cycle and DNA-repair genes at 35dpf, consistent with more cells undergoing meiosis.

A comparison of *amh* mutant juveniles to wild-type juveniles showed that numerous immune-related genes were up-regulated in mutants. These genes included interferon regulatory factor-7 (*irf7*), which is a paralog of the trout master sex-determining gene *sdY* that has been shown to be a duplicated, truncated interferon regulatory factor 9 (*irf9*) (Yano *et al*. 2012). The up-regulation of immune-related genes might reflect an inflammatory response to cell damage that accompanies the disruption of the development of Leydig cells and granulosa cells observed in histological sections and reflected in our *in situ* hybridization and transcriptome studies.

Transcriptomes of adult *amh* mutant testes differed substantially from those of adult wild-type testes. First, mutant testes up-regulated several genes that were greatly over-expressed in wild-type ovaries, likely reflecting the oocytes that histological sections revealed in *amh* mutant testes. These results show that Amh acts to block oocyte development in zebrafish testes. Second, adult *amh* mutant testes under-expressed most Leydig cell marker genes, consistent with our histology results and the finding that mouse *Amh* mutant males have disrupted Leydig cell development (Behringer *et al*. 1994). In contrast to the strongly altered Leydig cell markers in *amh* mutant transcriptomes, Sertoli cell marker genes were not strongly altered in *amh* mutants. We conclude that, although Amh is produced by and secreted from Sertoli cells, the lack of Amh function alters the developmental activities of Leydig cells, a conserved feature of Amh function. Thus, Leydig cells must express a receptor for Amh even though zebrafish lacks an ortholog of Amhr2, which helps form the Amh receptor in mammals and most fish. Third, *amh* mutant testes under-expressed testis-biased genes, consistent with the finding of ovotestes in *amh* mutants. Fourth, Despite *amh* mutant males accumulating substantial quantities of testis lobules, they greatly under-expressed genes involved in the production of mature sperm, verifying histology. The enormous testes in zebrafish *amh* mutants are consistent with zebrafish organ culture experiments that showed that Amh inhibits androgen-stimulated proliferation of spermatogonia (Skaar *et al*. 2011), thus, with less Amh, the inhibition should lessen, resulting in the accumulation of spermatogonia we observed. Together, our findings show that Amh signaling is required for normal development of Leydig cells, for the disappearance of ovotestes, for accelerating sperm maturation, and for the inhibition of spermatocyte proliferation.

Transcriptome analyses and in situ hybridization studies showed differences between adult ovaries in *amh* mutants and wild types, including the under-expression of marker genes for both granulosa cells and theca cells. In contrast, many oocyte marker genes were not differentially expressed, which is a bit surprising given the mis-expression of many of these genes in the putatively female groups (Groups21-1 and 35-1) in late larval and juvenile zebrafish. On the other hand, *amh* adult mutant ovaries over-expressed many genes involved in translation, as expected from the massive accumulation of young oocytes in mutant ovaries, and they under-expressed some ovary regulatory genes like *foxl2a, cyp19a1a,* and *gata4,* reflecting the loss of control of oocyte proliferation.

In tetrapods and most teleosts, Amh receptor type II (Amhr2) is expressed in both Leydig and Sertoli cells (Racine C *et al*. 199; Di Clemente N *et al*. 1994) and it mediates Amh signaling. Reference genomes of zebrafish and as we show, other cyprinids, however, have no ortholog of *Amhr2,* and gene loss is associated with a chromosome break at the ancestral site of the gene, thus, Amhr2 function is likely performed by another Bmpr2 paralog. In zebrafish, *bmpr2a* is over-expressed in adult wild-type testis vs. wild-type ovary and both *bmpr2a* and *bmpr2b* are under-expressed in *amh* mutant ovary vs. wild-type ovary, making these genes candidates for the elusive Amh receptor.

## ACKNOWLEDGEMENTS

We thank the University of Oregon Genomics & Cell Characterization Core Facility (Doug Turnbull and Maggie Weitzman) for their wonderful expertise, Clay Small for comments on the manuscript, and the NIH for support by grants R01 GM-085318 (J.H.P.), and IOS-1456737 (B.W.D.).

---

**Supplemental Figure S1.**
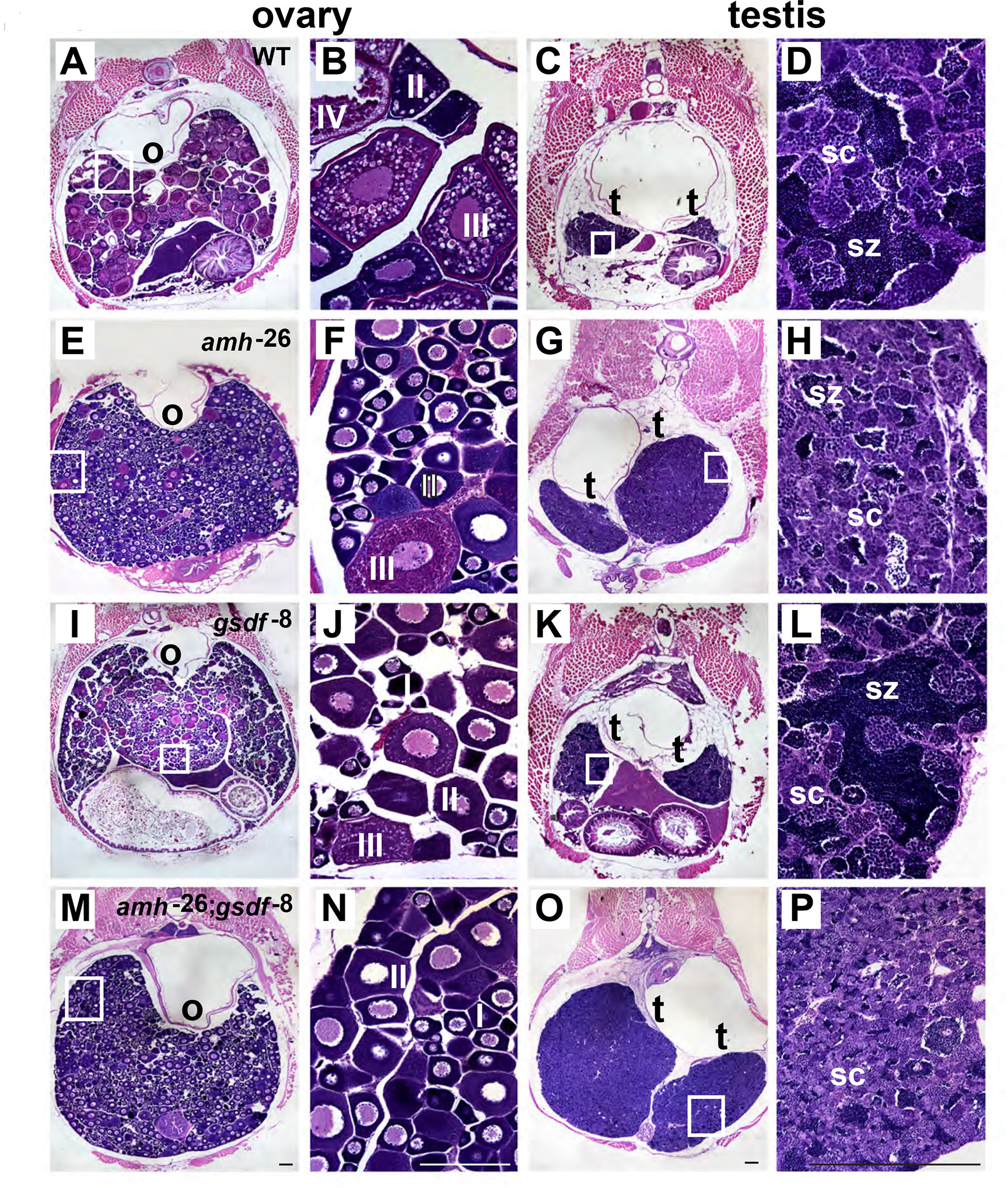
Mutant phenotypes for *amh;gsdf* double mutants. Cross-sections of gonads from one-year old adults. (A-D) Wild type (WT); (E-H) *amh^−^*^26^ mutants; (I-L) *gsdf* ^−8^ mutants; (M-P) *amh*^−26^;*gsdf* ^−8^ double mutants. (A, E, I, M) Ovary at low magnification. (B, F, J, N) High magnification of the boxed regions in A, E, I, M. (C, G, K, O) Testis at low magnification. (D, H, L, P) High magnification of the boxed regions in C, G, K, O. Abbreviations: I, II, III, IV: ovarian follicle stages 1 to 4; o, ovary; sc, spermatocytes; sz, spermatozoa; t, testis. Black scale bar in M for A, E, I and M; white scale bar in N for B, F, J and N; black scale bar in O for C, G, K and O; black scale bar in P for D, H, L and L. All scale bars: 100µm.

**Supplemental Figure S2.**
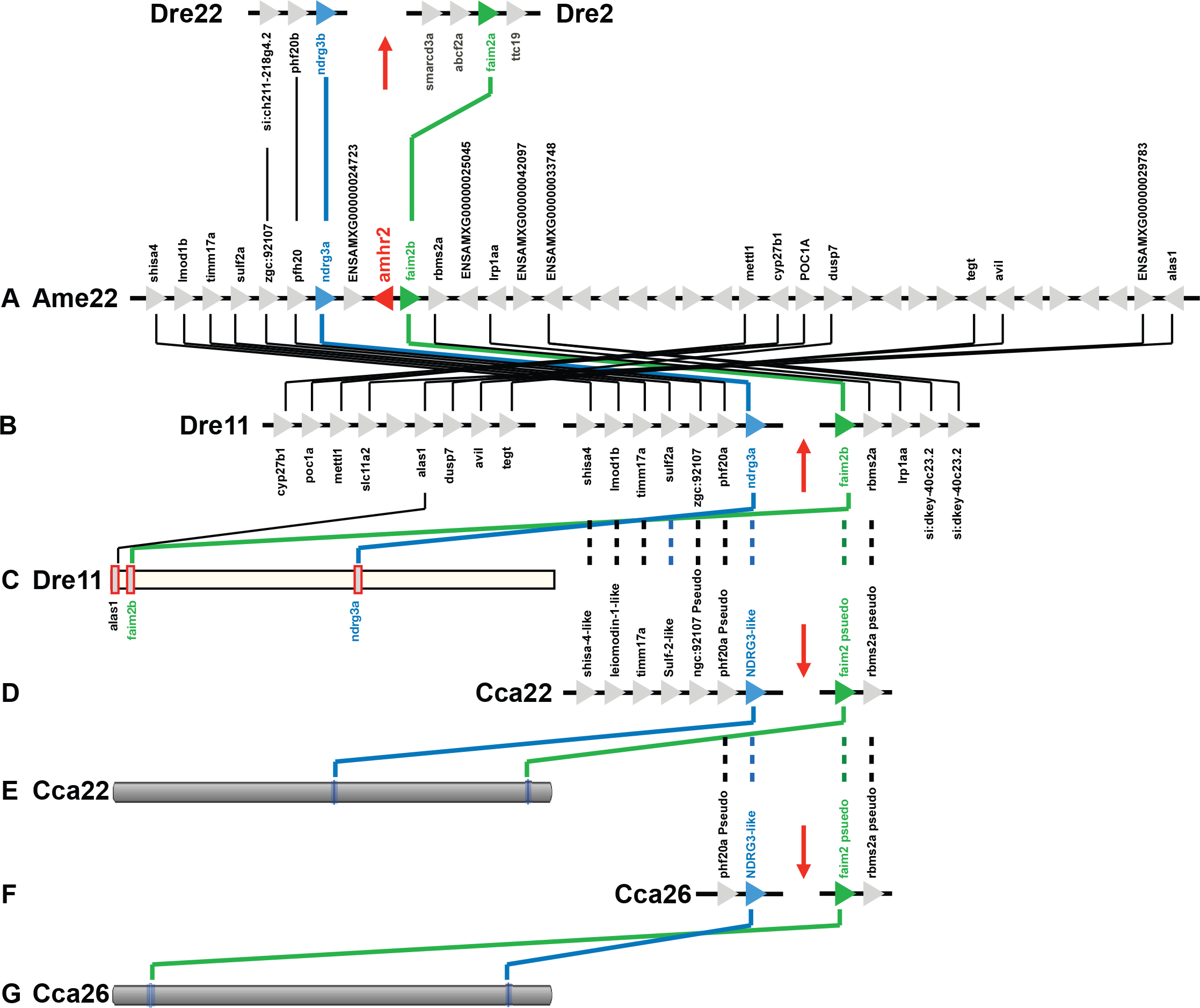
Conserved syntenies and loss of the zebrafish *amhr2* gene. A. A portion of chromosome 22 (Ame22) from the cavefish *Astyanax mexicanum* containing *amhr2*. B. Three portions of zebrafish chromosome 11 (Dre11) containing regions with conserved synteny to the *amhr2-*containing portion of the cavefish genome showing that the breakpoint of a chromosome rearrangement lies at the predicted location of *amhr2*. The portion of the figure above part A shows that paralogons of the zebrafish genome from Dre22 and Dre2 contain ohnologs derived from the teleost genome duplication and also lack *amhr2*. C. The three regions of the zebrafish genome shown in in part B occupy different positions along the entire chromosome Dre11. D. Portions of the common carp (*Cyprinus carpio*) genome on two parts of Cca22 with orthology to the *amhr2*-containing part of cavefish, which are broken at the expected site of the *amhr2* gene as in zebrafish. E. Positions of regions shown in part D on chromosome Cca22. F. The duplicated region from the carp genome duplication event co-orthologous to zebrafish chromosome Dre11. G. Positions of regions shown in part F on chromosome Cca22. Results show that *amhr2* loss occurred associated with a chromosome rearrangement breakpoint that is a shared feature of cypriniforms that occurred after cypriniform otophysans diverged from characiform and siluriform otophysans.

**Supplemental Figure S3.**
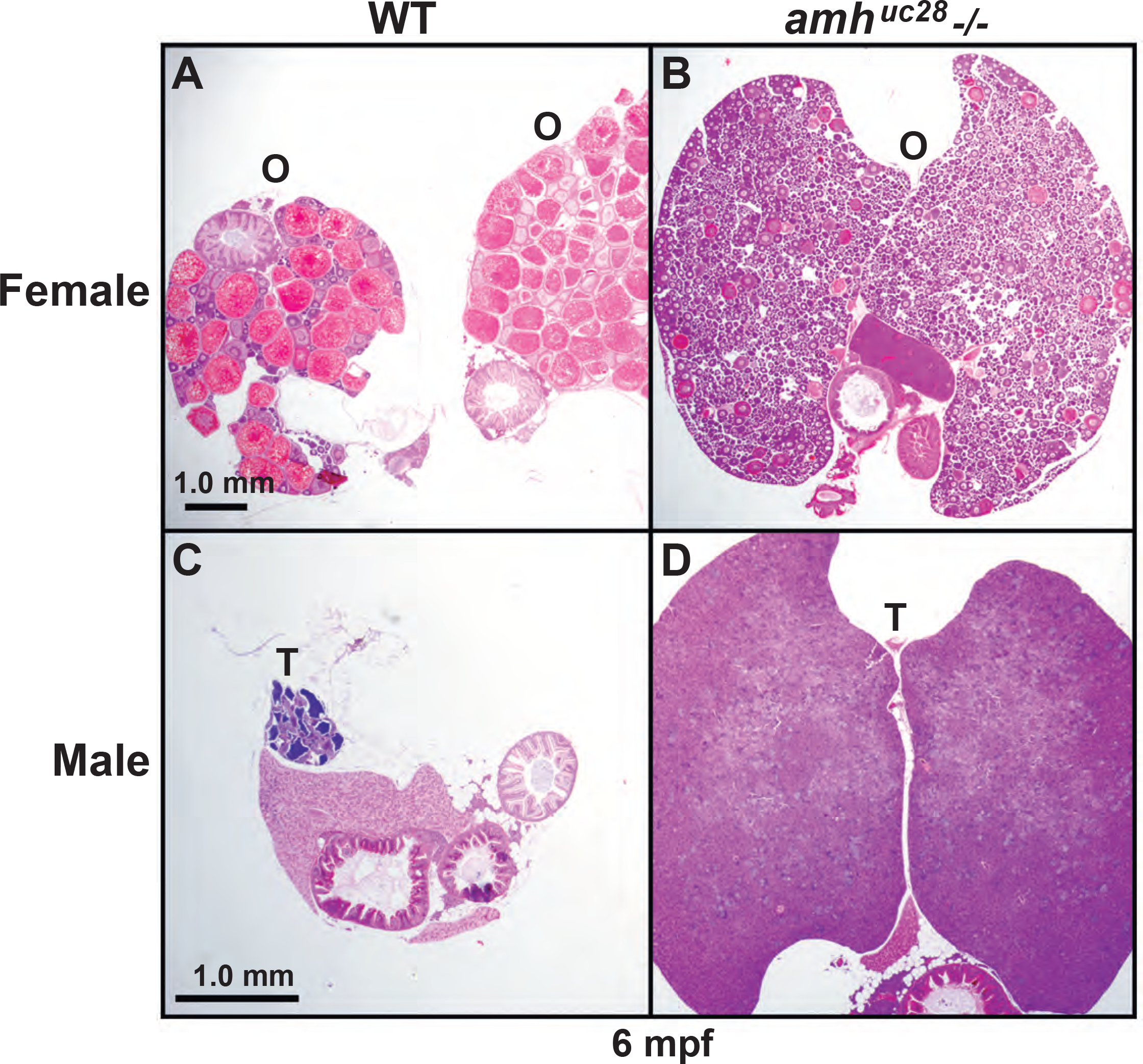
Gonadal phenotypes of the *amh^uc28^* allele visualized in hematoxylin and eosin stained histological sections of 6mpf animals. A. Wild-type (WT) female. B. *amh^uc28^* mutant female. C. Wild-type male. D. *amh^uc28^* mutant male. Abbreviations: O, ovary; T, testis. Scale bar: 1mm.

